# Incorporation of genome-bound cellular proteins into HIV-1 particles regulates viral infection

**DOI:** 10.1101/2023.06.14.544764

**Authors:** Manuel Garcia-Moreno, Robin Truman, Honglin Chen, Louisa Iselin, Caroline E. Lenz, Jeff Lee, Kate Dicker, Marko Noerenberg, Thibault J.M. Sohier, Natasha Palmalux, Aino I. Järvelin, Wael Kamel, Vincenzo Ruscica, Emiliano P. Ricci, Ilan Davis, Shabaz Mohammed, Alfredo Castello

## Abstract

The initial steps of the human immunodeficiency virus 1 (HIV-1) lifecycle are regulated by cellular RNA-binding proteins (RBPs). To understand the scope of these early host-virus interactions, we developed in virion RNA interactome capture (ivRIC), which allowed the comprehensive and systematic profiling of the proteins that interact with the HIV-1 genomic (g)RNA inside viral particles. ivRIC identified 104 cellular RBPs within the encapsidated HIV-1 ribonucleoprotein, many of which are typically found in the cellular nucleus. Notably, these nuclear RBPs interact with the HIV-1 RBP Rev, suggesting that they associate with HIV-1 gRNA during its nuclear life. Functional assays show that ivRBPs are important for HIV-1, including PURA and PURB, which control viral gene expression and infectivity through interaction with critical sequences in the gRNA. Our characterisation of the composition of the encapsidated ribonucleoprotein of HIV-1 uncovers new host-virus interactions that invokes new mechanisms for controlling HIV-1 infection.

## INTRODUCTION

Human immunodeficiency virus type 1 (HIV-1) RNA genome (gRNA) is reverse transcribed (RT) into DNA and integrated into the chromosome of the host cell. From this step onwards, cellular RNA-binding proteins (RBPs) become central players in the HIV-1 lifecycle, regulating or mediating virtually all molecular processes involving the viral RNA. Transcription and processing of HIV-1 RNAs is mediated by a plethora of cellular RBPs, including the RNA polymerase II, its associated co-factors and regulatory complexes, the capping and polyadenylation machinery and the spliceosome, amongst others^1^. HIV-1 produces several viral RNA species that can be divided in single spliced, fully spliced and unspliced (i.e. genome) RNAs^2^. Fully spliced HIV-1 RNAs are transported to the cytoplasm using the canonical mRNA export pathways involving many cellular RBPs^3^. However, unspliced and partially spliced RNAs are retained in the nucleus. The viral RBP Rev is expressed from a fully spliced RNA and imported into the nucleus, where it binds to the Rev response element (RRE)^4^ present in unspliced (gRNA) and single spliced HIV-1 RNAs. Rev then mediates the nuclear export of these RNAs through the CRM1 pathway, involving several cellular RBPs as co-factors^2^. Many RBPs are known to also contribute to the downstream steps of viral gene expression, including viral RNA stability and decay^5^, translation^6^, RNA transport, and viral particle formation^7^. Little is known about the potential involvement of cellular RBPs in subsequent processes of the HIV-1 lifecycle, such as viral particle formation and RT. However, recent discoveries suggest that RBPs may participate in these processes as well^7^.

It was originally thought that HIV-1 particles disassemble upon cell entry, releasing the genomic (g)RNA molecules into the cytoplasm for reverse transcription (RT). However, recent paradigm-shifting advances have challenged this view, showing that the capsid core remains intact during its transit through the cytosol and can be visualised traversing the nuclear pore and in the nucleoplasm^8, 9^. In addition, numerous lines of evidence revealed that RT can occur inside the capsid core, protecting the viral nucleic acids from the hostile cytosol^8, 10, 11^. Indeed, the capsid core harbours positively charged 8Å wide pores that allow deoxynucleotides to traverse from the cytosol to feed RT^10^. Interestingly, several individual examples of cellular RBPs regulating HIV-1 particle assembly^12^ and RT^13–20^ have been reported. However, the scope of cellular RBPs participating in these processes remain unknown. Moreover, we know very little about the mechanisms by which they are recruited to the gRNA and regulate viral particle formation and RT. Since the capsid core remains intact upon cell entry and RT occurs inside it, we hypothesise that relevant cellular RBPs should be packaged inside the HIV-1 particles in the producer cell and carried into the newly infected cell. In agreement, proteomic analyses of purified HIV-1 particles have revealed the presence of hundreds of RBPs within virions^7^. While exciting, these proteomic analyses were affected by biological and technical factors, including i) the uptake of a portion of the cytosol by budding particles, leading to the passive acquisition of bystander proteins; ii) the presence of extracellular vesicles with similar sizes to virions in the preparations; and iii) the lack of appropriate negative controls and/or quantitative information. Therefore, progress in understanding the RBP composition of virions requires the development of new strategies to differentiate between passive bystanders and functionally relevant proteins.

Here, we applied a new approach to determine the complement of cellular proteins that interact with the HIV-1 gRNA inside the viral particles. We discover that the *in virion* packaged genomic ribonucleoprotein (ivRNP) contains over one hundred cellular proteins, many of which are nuclear despite virion assembly occurring at the plasma membrane. To test if these nuclear RBPs associate with nuclear HIV-1 ribonucleoproteins (RBPs), we elucidated the first protein interactome of Rev in infected CD4+ T lymphocytic cells. To our surprise the ivRNP and the Rev interactome heavily overlap, suggesting a previously unknown connection between the nucleus and the formation of HIV-1 particles. Functional analysis revealed that the components of the ivRNPs play important regulatory roles in HIV-1 infection. For example, we found that PURA and PURB bind to critical regulatory elements on the HIV-1 gRNA and regulate viral gene expression and viral particle infectivity. Our study provides a new landscape of host-HIV interactions with regulatory potential.

## RESULTS

### ivRIC, a new approach to analyse the composition of the RNPs assembled into virions

Proteomic analysis of purified viral particles has revealed that hundreds of cellular proteins are incorporated into virions, including cellular RBPs^7^. Comparison of these datasets revealed sparse overlapping (Figure S1A), raising question about the biological significance of the identified proteins. The limited consistency of these studies is likely because total viral particle analysis is unable to discriminate between bystander proteins captured passively during virion assembly and RBPs actively interacting with the HIV-1 gRNA. To exclusively identify RBPs bound to gRNA in HIV-1 particles, we developed a new approach referred to here as *in virion* RNA interactome capture (ivRIC). In brief, viral particles are purified in a sucrose cushion, followed by protein-RNA ‘zero distance’ ultraviolet (UV) crosslinking, lysis under denaturing conditions and isolation of the polyadenylated HIV-1 gRNA with oligo(dT) magnetic beads via denaturing washes (Figure 1A). RBPs crosslinked to the gRNA are then released by RNase treatment and identified by quantitative proteomics. ivRIC was applied to HIV-_1mCherry-Nef_ particles (Figure S1B and S1C) purified from the supernatant of transfected HEK293T (Figure S1D) or infected CD4+ T lymphocytic cells (SupT1, Figure 1B). Interestingly, ivRIC showed that HIV-1 capsid (CA) is only isolated when forming part of the Gag polyprotein (also p55) and in a UV-dependent manner (Figure 1B and S1D). Free CA (also p24) was not detected in ivRIC eluates despite being very abundant (∼2k copies/virion) and proximal to the ivRNP. These results can be explained by the Gag’s RNA-binding properties, as it is the nucleocapsid (NC) polypeptide that interacts with RNA. Upon proteolytic processing of Gag-p55 by the HIV-1 protease (PR), NC remains attached to the gRNA whereas CA and matrix (MA) are released to form the inner capsid and outer matrix lattice (Figure 1C). In addition, the integrase (IN, also p31) was also enriched in ivRIC eluates in a UV-dependent fashion, which is consistent with its reported RNA-binding activity^21^ (Figure 1B). Conversely, the glycoprotein gp120 was present in inputs (i.e. viral particles) but absent in ivRIC eluates as it localises at the envelope, far away from the gRNA. Altogether, these results support the efficiency and selectivity of ivRIC at isolating viral proteins that are known to interact with the HIV-1 gRNA inside virions.

**Figure 1.**
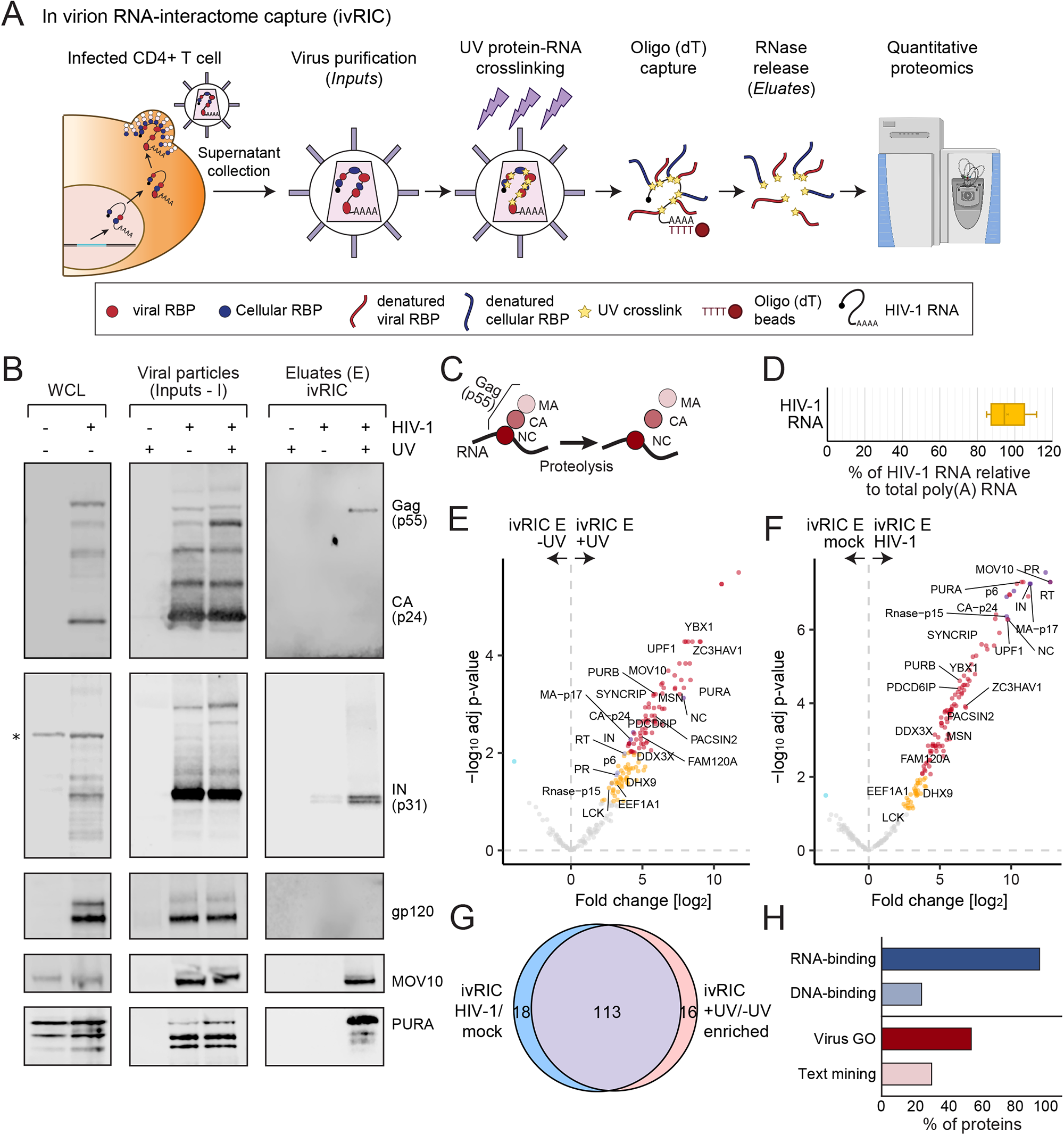
in virion RNA interactome capture (ivRIC) reveals the composition of the HIV-1 genomic ribonucleoprotein packaged in virions (ivRNP). A) Schematic representation of ivRIC. Samples and controls are depicted in Figure S1E. B) Western blotting of the whole cell lysates (WCL), inputs and eluates of a representative ivRIC experiment performed in CD4+ T-lymphocytic (SupT1) cells. Asterisk marks unspecific bands. C) Schematic representation of the proteolysis of the Gag polyprotein. MA, matrix; CA, capsid; NC, nucleocapsid. D) Relative proportion of the HIV-1 gRNA in eluates of ivRIC estimated by absolute RT-qPCR. E-F) ivRIC analysis of viral particles purified from the supernatant of SupT1 cells by quantitative proteomics. The volcano plots show the Log2 fold change and adjusted p-value of each protein detected in UV irradiated versus non-irradiated viral particles (E), or in the isolates from the supernatant of HIV-1-infected versus mock-infected cells (F). Red and dark blue dots are proteins enriched with 1% FDR, while orange and cyan dots are proteins enriched with 10% FDR. Grey dots are non-enriched proteins. G) Venn diagram showing the overlapping between the UV irradiated vs non-irradiated samples and HIV-1 infected versus mock-infected comparisons. H) Bar plots showing the proportion of ivRBPs in the ivRNP annotated with RNA- and DNA-binding (GO terms); virus-related (GO terms) and HIV-1-related (text-mining) functions. E, eluate.

The quality of ivRIC results relies on the efficient and specific isolation of HIV-1 gRNA. We thus quality controlled our results by quantifying the amount of viral RNA in ivRIC eluates. Notably, HIV-1 gRNA represented ∼95% of the isolated RNA (Figure 1D), which agrees well with the previously described proportion of viral versus cellular polyadenylated RNA found in purified HIV-1 particle preparations^22^. The dominance of HIV-1 gRNA in eluates supports the high efficiency and selectivity of ivRIC, while implying that viral RNA must be the major (if not the sole) contributor to the subsequent proteomic analyses.

### Uncovering the composition of the ivRNP

To comprehensively and systematically discover the cellular proteins that compose the HIV-1 ivRNP, we analysed the eluates of ivRIC from CD4+ T lymphocytic cells by quantitative proteomics. We included several controls to determine if protein identification relies on i) UV crosslinking (+UV vs -UV), ii) HIV-1 infection (infected vs mock), and iii) protein abundance (whole cell proteome [WCP] and total viral particle proteome [inputs]) (Figure S1E and F). Principal component analysis revealed that eluates from UV-irradiated HIV-1-infected cells strongly differed from control samples in which UV crosslinking or infection was omitted (Figure S2A). 120 unique cellular and 9 viral proteins were statistically enriched in ivRIC eluates in a UV-dependent manner (Figure 1E), with nucleocapsid (NC, p11) exhibiting the highest fold change of all HIV-1 polypeptides (256-fold enrichment). This agrees with the notion that NC tethers Gag/p55 and Gag-Pol polyproteins to the viral RNA. 122 cellular and 9 viral polypeptides were enriched in samples from HIV-1-infected cells over mock-infected controls (Figure 1F). Importantly, 104 cellular and the 9 viral proteins (113 in total) were consistently enriched over the two controls (Figure 1G), reflecting that these proteins i) interact with the gRNA isolated from viral particle preparations, and ii) are not derived from cellular vesicles with virus-like sizes. We refer to these high confidence components of the ivRNP as *in virion* RNA-binding proteins (ivRBPs).

We next validated our proteomic results using Western blot as an orthogonal approach. Beyond the excellent recovery of viral proteins known to interact with RNA (see above), the cellular helicase MOV10 was strongly enriched by ivRIC in a UV and infection-dependent manner (Figure 1B), reinforcing earlier findings showing its incorporation into HIV-1 particles^20, 23, 24^. Our results further show that MOV10 actively engages with gRNA in the context of the ivRNP. Moreover, other ivRBPs such as PURA, CIRBP and SUB1 were also detected in ivRIC eluates by Western blotting in a UV and infection dependent manner (Figure 1F and Figure S2B). Collectively, these results confirm the ability of ivRIC to uncover cellular RBPs that interact with HIV-1 gRNA inside virions.

To gain functional insights into the composition of the HIV-1 ivRNP, we used available literature and gene ontology (GO) annotation. As expected from a high-quality RNA interactome, most ivRBPs were annotated with the ‘RNA binding’ GO term and were linked to functions associated with RNA metabolism (Figure 1H, 2A and S2C). Interestingly, ∼25% of the ivRBPs also contain DNA-binding activity (Figure 1H and 2A), which is potentially very significant given the RNA/DNA duality of the HIV-1 genome^10, 25^. Moreover, over half of the discovered ivRBPs are annotated with GO terms associated with virus infection, and ∼25% of them also linked to HIV-1 (Figure 1H). Indeed DHX9^16, 17^, EEF1A^18^, UPF1^14^, MOV10^20, 24^, PDCD6IP (ALIX)^12, 26^, PACSIN2^27^ and LCK^28^ have been related to viral particle assembly or infectivity. Other proteins such as DDX3^29^, YBX1^30^, PURA^31^ and ZC3HAV1 (ZAP)^32, 33^ have been associated with the HIV-1 lifecycle, but their roles within viral particles (if any) are still unknown. Moreover, dozens of ivRBPs have no prior connection with HIV-1, calling for further investigation.

**Figure 2.**
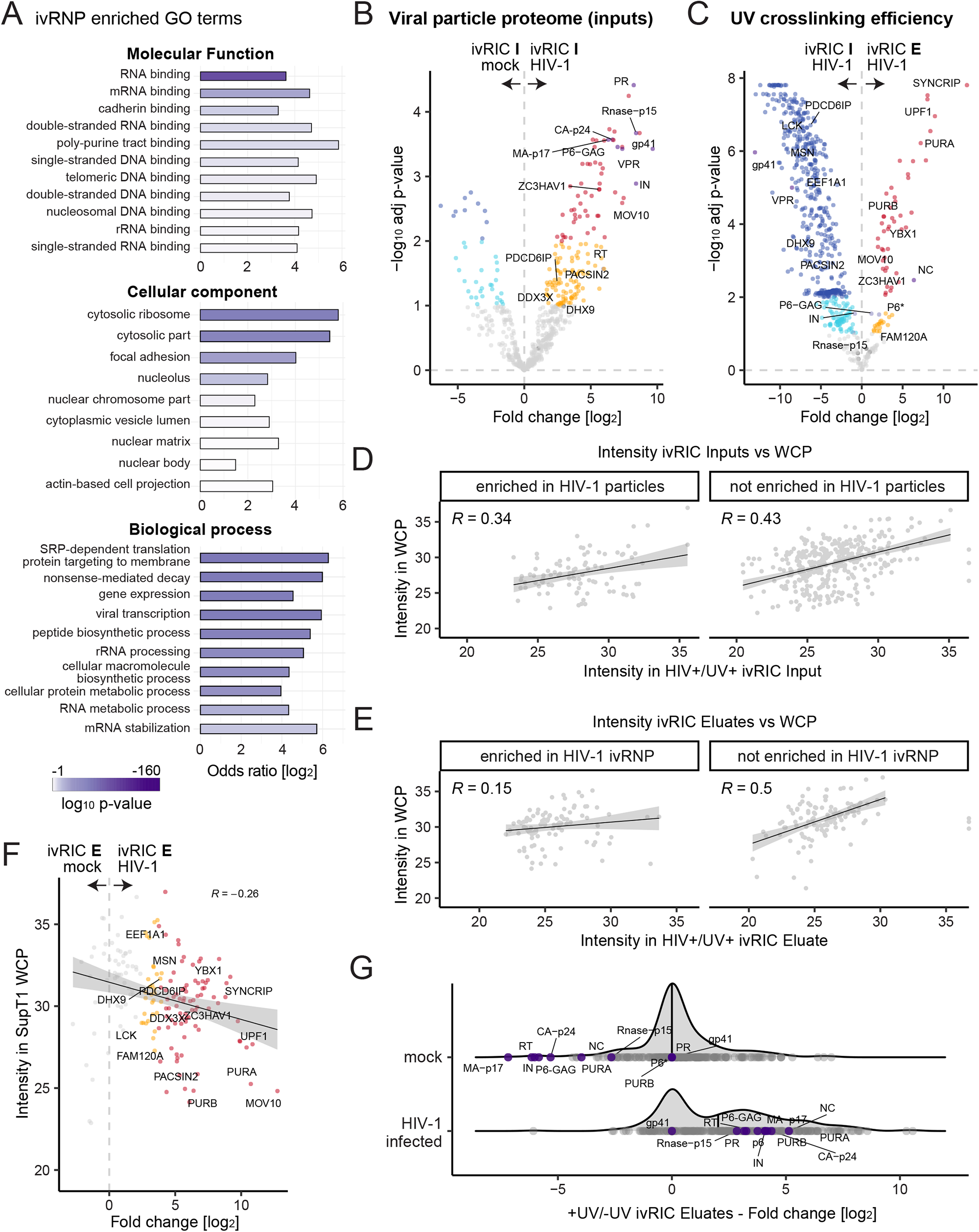
Analysis of the properties of ivRBPs. A) Analysis of the GO terms enriched in the ivRNP over the cellular proteome. B-C) Volcano plots showing the Log2 fold change and adjusted p-values for each protein (dot) in the comparison ivRIC inputs (I, purified viral particles) from HIV-1 infected and mock-infected cells (B) and ivRIC inputs versus eluates (E, ivRNP) (C). Red and dark blue dots are proteins enriched with 1% FDR, while orange and cyan dots are proteins enriched with 10% FDR. Grey dots are non-enriched proteins. D-E) Scatter plots showing the intensity of proteins in ivRIC inputs (D) or eluates (E) against the whole cell proteome (WCP). F) Scatter plot showing the protein intensity in the WCP (y axis) and the Log2 fold change in the ivRIC eluates from mock and HIV-1 infected as in (B-C). G) Density plots showing the distribution of fold changes in UV irradiated and non-irradiated samples for proteins detected in HIV-1 infected and mock supernatants. Cellular proteins and viral proteins are shown in grey and purple, respectively.

### The total proteome of HIV-1 particles provides new insights into the RNA-binding properties of ivRBPs

To determine the composition of the HIV-1 particles prior to gRNA isolation, we analysed the inputs of the ivRIC experiment by quantitative proteomics (Figure 1A and S1E). 187 proteins were significantly enriched over mock controls, representing the total viral particle proteome (Figure 2B, S3A-B). In contrast to the HIV-1 ivRNP, the proteome of viral particles was enriched in GO terms and pathways related to the plasma membrane and immunological receptors, likely reflecting the envelope acquired during budding from the plasma membrane (Figure S3C-E). ‘RNA binding’ was also enriched, but the odds ratio and p-value were substantially lower than those of the HIV-1 ivRNP (Figure S3C and E). The different prevalence of GO terms between the two datasets is expected as the ivRNP is a subset of the total HIV-1 particle.

RBPs can display different modes of RNA binding, ranging from universal to very selective, and from long-lived to transitory RNA binding. The RNA-binding properties of an RBP do not only impact its function (i.e. global vs specific and transitory vs stable)^34^, but also its ability to crosslink to RNA. Indeed, UV-induced, RNA-to-protein crosslinking is enhanced by long-lived and geometrically optimal interactions between nucleotides and amino acids^35^. To assess the binding properties of the ivRBPs, we compared the protein intensity in eluates (ivRNP) and inputs (viral particles) of the ivRIC experiment. This analysis revealed two groups of proteins (Figure 2C): i) ivRBPs with high eluate/input protein intensity ratios (i.e. high ‘crosslink-ability’), and ii) ivRBPs with low eluate/input ratio (i.e. low crosslink-ability). ivRBPs in the first group are expected to establish long-lived and geometrically optimal interactions with the gRNA, while the opposite is expected for the second group. Although strong and specific interactions are expected to be functionally relevant, there are many examples of transitory and low affinity interactions with fundamental roles in RNA metabolism^34^. Indeed, some of the ivRBPs with low crosslink-ability to HIV-1 gRNA, such as PDCD6IP^12, 26^, LCK^28^, DHX9^16^, have been shown to play important regulatory roles in viral particle formation and RT.

### ivRBPs are incorporated selectively into viral particles

To discriminate between abundance-driven passive incorporation of bystander proteins and selective packaging of proteins into virions, we normalised the protein intensity in ivRIC inputs (viral particles) and eluates (ivRNP) against protein intensity in the whole CD4+ T cell proteome (WCP). Interestingly, the intensity of proteins in inputs correlated well with protein abundance (R=∼0.4; Figure 2D), suggesting that protein levels do influence overall virion composition. By contrast, protein intensity correlation was substantially lower when comparing the proteins enriched in the ivRNP with the WCP (R=0.15) (Figure 2E). Non-enriched proteins exhibited similar trends to inputs (R=0.5). When considering the fold change instead of raw intensity, the result was even more clear, with ivRBPs displaying anticorrelation with the WCP (Figure 2F and S2D). This result indicates that the stronger a protein is enriched in the ivRNP, the lower its abundance in the cell. Together, these data strongly favour the notion that ivRBPs are selectively incorporated into viral particles.

A remaining question is whether ivRBPs are present in sufficient quantities in ivRNPs to have functional consequences in infection. To test this, we compared the protein intensity and fold change distribution in eluates of ivRIC from mock- and HIV-infected cells. Interestingly, many cellular ivRBPs exhibited similar intensity levels and fold changes to viral proteins with high stoichiometry in viral particles (Figure 2G, grey vs purple). Several known regulators of HIV-1 infectivity were amongst the most prevalent ivRBPs in virions, including UPF1^14^ and MOV10^23^. However, other prevalent ivRBPs such as PURA, PURB, and ZC3HAV1/ZAP have no known roles inside HIV-1 particles. The high abundance of MOV10 and PURA in particles was confirmed using Western blots (Figure 1B). Our results are thus compatible with the notion that ivRBPs play potential roles in infection.

### Elucidating the Rev interactome in infected CD4+ T cells

The presence of many nuclear RBPs (Figure 2A) in the ivRNP is surprising as HIV-1 particle assembly occurs at the plasma membrane. We hypothesised that these proteins may associate with the gRNA in the nucleus and remain bound to it until its packaging into viral particles. To test our hypothesis, we focused on a key component of the HIV-1 nuclear genomic ribonucleoprotein (gRNP), called Rev^2^. Therefore, the protein interactome of Rev is expected to reflect the composition of the gRNP to a high extent. There have been several attempts at establishing the Rev interactome, however, these approaches were limited because Rev was expressed in non-infected cells and/or using cell types that are not infected by HIV-1, leading to very poor overlap between datasets (Figure S4A)^2, 36–38^. To study the Rev interactome in infected CD4+ T lymphocytic cells, we generated a chimeric HIV-1 replicon that expressed Rev fused to Flag-Myc (HIV-1_-R-E-Rev-Flag-Myc_) or HaLo tag (HIV-1_-R-E-Rev-HaLo_) (Figure 3A and S4B).

**Figure 3.**
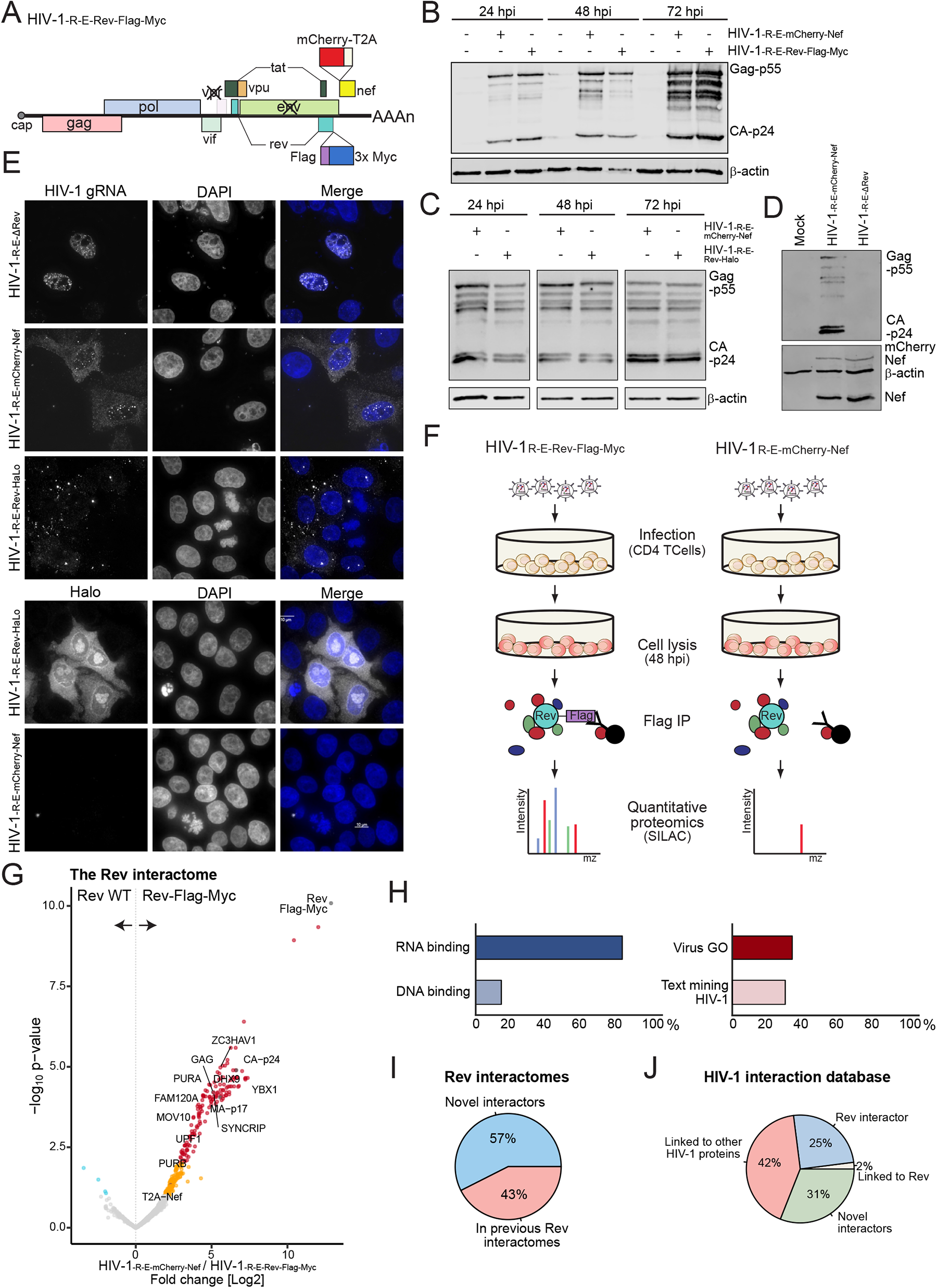
The Rev protein-protein interactome. A) Schematic representation of the HIV-1-R-E-Rev-Flag-Myc construct. B-D) Western blotting showing viral protein expression in CD4+ T lymphocytic cells (SupT1) infected with HIV-1-R-E-Rev-Flag-Myc (B), HIV-1-R-E-Rev-Halo (C), HIV-1-R-E-ΔRev (D) and control HIV-1-R-E-mCherry-Nef. hpi, hour post infection. E) Fluorescence microscopy analysis of HeLa cells infected with the indicated HIV-1 construct. Viral RNA was detected by single molecule in situ hybridisation (smFISH) (upper panels), Rev-HaLo with fluorescent HaLo reagents (bottom panels), and the nuclei was stained with DAPI. F) Schematic representation of the Rev protein-protein interaction experiment, using chimeric pseudotyped viruses (HIV-1-R-E-Rev-Flag-Myc and HIV-1-R-E-mCherry-Nef as negative control) to infect CD4+ T lymphocytic cells (SupT1). G) Volcano plot showing the Log2 fold change and adjusted p-values for each protein (dot) comparing the eluates of the Rev-Flag-Myc immunoprecipitation (IP) versus the untagged control (Rev WT). Red and dark blue dots are proteins enriched with 1% FDR, while orange and cyan dots are proteins enriched with 10% FDR. Grey dots are non-enriched proteins. H) Bar plots showing the proportion of Rev interactors annotated with RNA- and DNA-binding (GO terms); virus-related (GO terms) and HIV-1-related (text-mining) functions. I) Proportion of novel and previously identified Rev interactors. J) Pie chart showing the proportion of proteins classified as novel Rev interactors and previous annotations in HIV-1 NCBI database.

To assess if tagging affects Rev function, we generated HIV-1 particles by co-transfection of the replicons with the glycoprotein of the vesicular stomatitis virus (VSV G). Infection of SupT1 cells with these viruses led to normal levels of Gag/p55, processed CA/p24 and mCherry-Nef, when compared to the parental HIV-1_-R-E-mCherry-Nef_ (Figure 3B-C and S4C). By contrast, nullification of Rev by adding a stop codon (HIV-1_-R-E-ΔRev_) led to undetectable levels of Gag/p55 (Figure 3D). Single molecule *in situ* RNA hybridisation (smFISH) of cells infected with HIV-1_-R-E-Rev-HaLo_ showed that the expression of the tagged Rev overcomes the nuclear accumulation of gRNA observed in absence of Rev (Figure 3E). Rev exhibits a nucleolar localisation when overexpressed in non-infected cells^39^. Conversely, expression of Rev from our construct led to additional localisation in nuclear pores and nucleoplasm (Figure 3E). Presence of Rev at nuclear pores is compatible with its known role in HIV-1 RNA nuclear/cytoplasmic export^2^. Collectively, our results confirmed that Rev is functional when tagged with Flag-Myc or HaLo.

To elucidate the Rev interactome, we infected SupT1 cells with the chimeric viruses, followed by Flag immunoprecipitation (IP) at 48 hours post infection (hpi) in presence of RNases (Figure 3F). Proteomic analysis revealed high sample correlation for Rev-Flag IPs (Average Pearson correlation, R= 0.85; Figure S4D-E), and that Rev was the most enriched protein in the IP (Figure 3G and S4F). In addition to Rev, 284 cellular proteins were significantly enriched in IP eluates at 10% false discovery rate (FDR) (Figure 3G). 81% of the interactors were RBPs themselves (Figure 3H and S4G), reflecting Rev’s prominent role in viral RNA metabolism. We noticed the presence of large number of proteins from the ribosome and spliceosome in the IP eluates (Figure S4H), supporting previous finding that Rev is a regulator of HIV-1 RNA splicing and translation^2^. In addition, Rev interacts with complexes involved in chromatin function and RNA polymerase II transcription (Figure S4H), suggesting that it may engage with viral RNA co-transcriptionally.

The previously established Rev interactomes showed very poor overlapping between them, raising questions over their biological significance^2^. However, 43% of the proteins identified here were also reported in these earlier studies (Figure 3I). Our results thus reconcile the contradictions in these datasets, while still adding many additional Rev interactors. We also observed a strong overlap between our Rev interactome and the annotation in the NCBI HIV database, with 25% of the proteins being already annotated as Rev interactors and a further 42% previously linked to HIV-1 in some capacity (Figure 3J). Altogether, these results highlight the consistency of our Rev interactome with previous data, while adding new Rev-host protein interactions with unknown roles in HIV-1 infection.

### The HIV-1 ivRNP heavily overlaps with the Rev protein-protein interactome

Nuclear RBPs present in the ivRNP may associate with the HIV-1 gRNA in the nucleus of the infected cell. To determine to what extent the ivRNP resembles to the nuclear gRNP, we compared the ivRIC (Figure 1) and Rev IP results (Figure 3). Surprisingly, 68% of the components of the ivRNP are also interactors of Rev (Figure 4A). Indeed, we observed that the proteins identified in both the Rev interactome and the ivRNP displayed similar fold changes in both datasets, showing a striking Pearson correlation of >0.61 (Figure 4B). This suggests that the stoichiometry of these proteins in the Rev interactome and ivRNP is likely similar. Proteins shared between these two datasets are mostly involved in RNA transcription, splicing, nuclear export, and translation (Figure 4C). By contrast, proteins only present in the ivRNP are associated mainly with membrane biology, including membrane trafficking, endocytosis, and vesicle transport. Our results are thus compatible with two populations of ivRBPs: one that associates with the gRNA in the nucleus and remains associated with it, and another that engages with the gRNA later at the plasma membrane. By contrast, proteins specific to the Rev interactome are nuclear and are broadly involved in chromosome biology, DNA damage, Tat-mediated transcription, rRNA processing and 3’ to 5’ RNA decay amongst others (Figure 4C). Interestingly, we also observed that the previously established Gag^40^ and Staufen^41^ interactomes (proxies for the cytoplasmic gRNPs) overlap to some extent with the ivRNP and the Rev interactome (Figure S5A). Moreover, several ivRBPs and Rev interactors were previously found associated with HIV-1 RNA in infected cells (Figure S5B)^42, 43^. These findings strongly suggest that a wide set of RBPs engage with HIV-1 gRNA during the nuclear and cytosolic life of the gRNA and remain bound to it during the formation of the viral particles. The striking functional connection between nuclear proteins and plasma membrane-assembled virions warrants further investigation.

**Figure 4.**
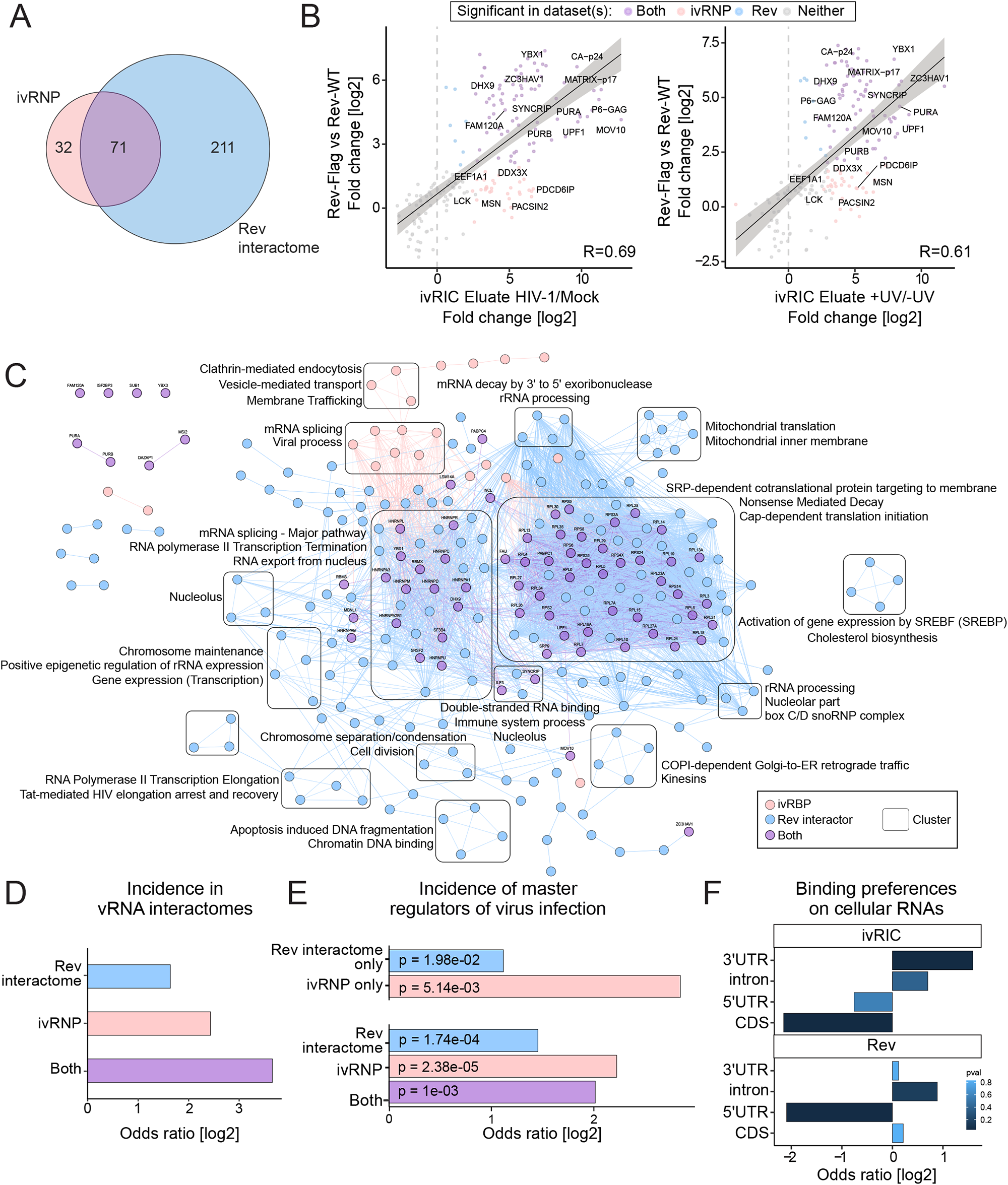
Comparison of the ivRNP and the Rev protein-protein interactome. A) Venn diagram showing the overlapping between the ivRIC (ivRNP) and the Rev IP (Rev interactome) experiments. B) Scatter plots comparing the Log2 fold change of the Rev IP (y-axis) and the ivRIC eluates (x-axis) against their respective controls. C) Pairwise STRING network comparison of the ivRNP and the Rev interactome indicating consistencies and divergences between the two datasets. Clusters are boxed and their associated top GO enrichment terms are shown. D) Odds ratio of proteins in ivRNP and/or Rev interactome that are identified in the RNA interactomes of other viruses (dataset from 44). E) Odds ratio of proteins reported to cause phenotypes in virus infection in the siRNA and CRISPR/cas9 screenings collected in 45. We consider a ‘master regulatory’ protein to any candidate whose perturbation causes phenotypes in viruses from three or more families. F) Analysis of the binding preferences of ivRBPs and Rev interactors using the eCLIP database (ENCODE)46. Odds ratio of binding sites in 5’ UTRs, CDSs, introns and 3’ UTRs was estimated for each protein in the ivRNP and Rev interactome and the overall preferences of each protein group was represented as a bar plot.

To test whether RBPs in the ivRNP and Rev interactome are specific to HIV-1 gRNA or have widespread binding activity across viral RNAs, we collected all the viral RNA interactomes available to date comprising 21 datasets across 11 viruses^44^. Our results show that the ivRNP and the Rev interactome are both enriched in proteins with widespread binding to RNAs from different viruses (Figure 4D and S5C). To assess if this broad-spectrum viral RNA-binding activity is linked to widespread regulatory activity in virus infection, we compiled all siRNA and CRISPR-cas9 screenings of viral fitness generated to date, which include 36 viruses and 18 viral families^45^. Interestingly, ∼20% and ∼13% of the components of the ivRNP and Rev interactome that were tested in the screenings caused phenotypes in at least three different viral families (Figure 4E and S5D).

The observed binding of ivRBPs and Rev interactors to the RNAs of different virus species, together with their potential master-regulatory roles in infection, could be derived from low binding specificity that enable them to engage with a wide range of RNA sequences. To test this, we examined the RNA-binding preferences of all the RBPs in these two datasets for whose enhanced crosslinking and immunoprecipitation (eCLIP) sequencing data is available in ENCODE^46^. Most of the proteins tested exhibited sequence-specificity as well as preferences for defined regions on cellular RNAs (Figure 4F and S5E-F). Despite a large overlap between the ivRNP and the Rev interactome, we observed some differences in the overall binding preferences, with 3’ UTRs and coding sequences (CDS) being predominant binding sites for ivRBPs, and introns and 5’ UTR for Rev interactors (Figure 4F). This divergence can be explained by the fact that the Rev interactome is substantially larger and include additional RBPs, many of which are involved in nuclear processes (Figure 4A and C). Our results therefore revealed binding preferences across the ivRNP and the Rev interactome components that are compatible with specific RNA binding.

### PURA and PURB are regulators of HIV-1 infection

To test if ivRBPs and Rev interactors are functionally relevant to HIV-1 infection, we generated knock out CD4^+^ T lymphocytic cells for several candidates. These included the transcriptional activator protein Pur-alpha (PURA) (Figure 1B), its homologue Pur-beta (PURB) and FAM120A that were identified by both ivRIC and Rev protein-protein interaction analysis. We also selected moesin (MSN) that was present only in the ivRNP, as expected from a membrane-associated protein. KOs were generated using Nanoblades^47^, leading to cells lacking PURA, PURB, FAM120A or MSN that had no detectable effects in cell viability, proliferation or cytotoxicity (Figure S6A-D).

To test the effects of these proteins on HIV-1 fitness, we monitored HIV-_1mCherry-Nef_ infection of CD4+ T lymphocytic WT and KO cells by flow cytometry (Figure 5A). Interestingly, all KO cell lines except for FAM120A exhibited a substantially lower number of mCherry-positive cells (Figure 5B and S6E), suggesting that PURA, PURB, and MSN are important for viral gene expression and/or spread in CD4+ T lymphocytic cells. The effects on HIV-1 gene expression were confirmed for PURA and PURB KO cells by Western blot, showing reduced levels of Gag, Gag-Pol and their proteolytic products (Figure 5C and S6F-G).

**Figure 5.**
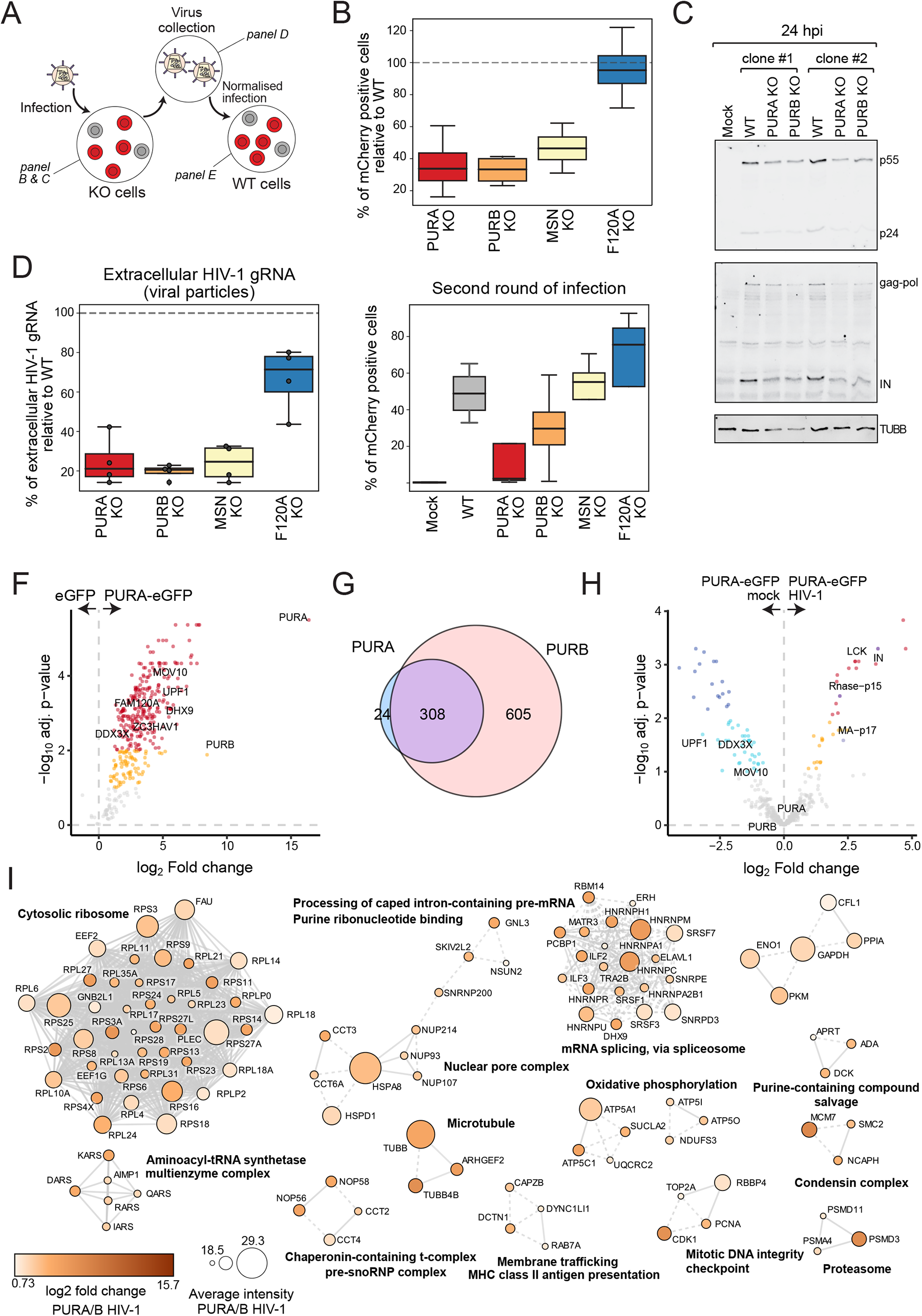
PURA and PURB regulate HIV-1 gene expression and infectivity. A) Representation of the experimental design. B) Flow cytometry analysis of WT and KO SupT1 cells infected with HIV-1mCherry-Nef for 48h. Y-axis show the percentage of KO cells expressing mCherry as compared to WT cells (dashed line). C) Western blotting analysis of WT, PURA and PURB KO SupT1 cells infected with HIV-1mCherry-Nef for 24 h. See quantification in Figure S6G. D) RT-qPCR analysis of the HIV-1 gRNA in the supernatant of WT and KO infected SupT1 cells from panel B. E) Viral particles purified in (D) from KO and WT cells were used to infect WT SupT1 cells upon normalisation by gRNA levels. mCherry positive cells were quantified by flow cytometry at 48 hpi. F) Volcano plot showing the proteins enriched in PURA-eGFP IP over the eGFP control IP Jurkat Flp-In T-REx cells expressing these proteins. Dots represent proteins enriched with 1% FDR (red and dark blue) or with 10% FDR (orange and cyan). Grey dots are non-enriched proteins. G) Venn diagram comparing the proteins that are enriched in both PURA-eGFP and PURB-eGFP IPs. H) Volcano plot showing the proteins enriched in mock versus HIV-1 infected cells. Colour code as in F. I) STRING clustered network showing physical (solid lines) and functional (dashed lines) connections between interactors for both PURA-eGFP and PURB-eGFP. Top GO enriched terms are shown for each complex.

To assess if viral production is affected by the lack of these ivRBPs, the supernatant of KO and WT cells was collected and analysed by RT-qPCR (Figure 5A and D). We observed a remarkable decrease of HIV-1 gRNA in the supernatant of KO cells, which correlated well with the reduced infection observed by flow cytometry (Figure 5B vs D). These results confirmed the importance of PURA, PURB, MSN and FAM120A in HIV-1 infection by an orthogonal approach.

The identification of PURA, PURB, MSN and FAM120A in the ivRNP suggests that they may also play a role in virus infectivity. To assess this, we infected WT cells with the same number of viral particles generated in WT or KO cells, using the RT-qPCR data for normalisation (Figure 5A and D). Viruses produced in MSN KO cells exhibited no differences in infectivity when compared to those generated in WT cells, while a slight increase was observed for FAM120A KO samples (Figure 5D). By contrast, viruses generated in PURA and PURB KO displayed a substantial decrease in HIV-1 positive cells, with PURA deficient viruses having the lowest infectivity (Figure 5E). Altogether, our results reveal that all tested ivRBPs participate in HIV-1 gene expression, while PURA and PURB have additional roles in the infectivity of viral particles.

### The PURA and PURB interactome in HIV-1 infected CD4+ T lymphocytic cells

To further characterise PURA and PURB roles in infection, we carried out an IP and proteomics experiment to determine their protein-protein interactomes. To achieve this in a relevant cell line, we generated Jurkat Flp-In T-REx cells to enable expression of PURA and PURB fused to eGFP in a CD4+ T lymphocytic line in a doxycycline-induced dependent manner (Tet-on). Expression of PURA-eGFP and PURB-eGFP was confirmed by Western blotting and fluorescence microscopy (Figure S7A and B). Induced cells were infected with HIV-_1mCherry-Nef_ and expression of both green and red fluorescent proteins was confirmed by flow cytometry (Figure S7C). At 48h, mock and infected cells were lysed and the eGFP fusion proteins were IPed with GFP nanobodies (GFP_Trap). The specific enrichment of these proteins in eluates was confirmed by silver staining and Western blotting (Figure S7D and E). Proteomic analysis of the eluates revealed hundreds of proteins that specifically co-precipitate with PURA-eGFP and PURB-eGFP (Figure 5F and S7F). Both datasets largely overlap, with 93% of the PURA interactors being also present in PURB IP (Figure 5G), suggesting that their functions are either linked or mutually exchangeable. PURB-eGFP IP dataset was larger, probably due to the higher expression of this protein when compared to PURA-eGFP (Figure S7B, D and E). While we observed differences in overall protein intensity (Figure S7G), the two IPs correlated well, suggesting that the composition of eluates was very similar (Figure S7H). Interestingly, PURB was the protein with highest fold change in PURA-eGFP IP and *vice versa* (Figure 5F and S7F), suggesting that both proteins associate in heteromeric complexes. Notably, Gag and Gag-Pol components such as MA, IN, RT, and the RNase H p15 were highly enriched in PURA-eGFP and PURB-eGFP IPs. Moreover, PURA and PURB interact with several cellular pathways and components related to cellular gene expression, including the condensin complex, the spliceosome, the nuclear pore, and ribosome (Figure 5I). These results suggest that PURA and PURB participate in several steps of the life cycle of cellular and HIV-1 RNAs. Interestingly, several microtubule-associated and membrane trafficking proteins were enriched in the pull down, indicating that PURA and PURB might also be involved in the transport of viral RNPs across the cell. A wide range of proteins exhibited differential association with PURA and PURB upon HIV-1 infection, many of which are involved in splicing and translation (Figure 5H, S7F and I). This included a few ivRBPs such as MOV10, that displayed a consistent decrease in PURA/B-binding upon infection. Altogether, these results highlight the engagement of PURA and PURB with a wide range of cellular machineries that are important for the life cycle of HIV-1 RNAs. Moreover, they also interact with Gag and Gag-Pol components that are essential for the formation of HIV-1 particles and early steps of viral infection.

### PURA and PURB bind to key regulatory regions in cellular mRNAs

We next aimed to elucidate the RNA-binding preferences of PURA and PURB. For this, we applied iCLIP2^48^ to HIV-1 infected and uninfected PURA-eGFP and PURB-eGFP Jurkat cells (Figure S7C). iCLIP2 employs UV cross-linking, RNase treatment, IP, cDNA library generation and sequencing to determine the footprints of RBPs on their target RNAs in a genome-wide manner and with single nucleotide resolution. Both PURA-eGFP and PURB-eGFP were strongly enriched in IP eluates (Figure S8A and B). Moreover, ligation of a fluorescently labelled DNA adapter to the co-precipitated RNA revealed a high molecular weight smear that is compatible with PURA/B-RNA complexes (Figure S8C). Upon cDNA generation and PCR amplification, the sequencing of the libraries revealed that PURA and PURB interact predominantly with protein coding cellular RNAs (i.e. mRNAs), with a bias towards coding sequences (CDS) followed by 3’ UTRs (Figure 6A and B and S8D-E). Binding sites in CDS, 5’ UTRs and introns displayed intriguing distributions. 5’UTR binding sites were prominent at the beginning (near the cap) and end (translation start codon) (Figure 6C). Similarly, the binding site density at CDS displayed two notable peaks near the translation start and stop codons (Figure 6C). The binding sites in introns also had a dual distribution ramping up at the 5’ and 3’ splice sites (Figure 6D). No bias was observed for 3’ UTRs. The striking binding site accumulation at the beginning and end of 5’ UTRs, CDS and introns overlapped or was proximal to key regulatory elements such as the cap, the start and stop codons and the splice sites, suggesting that PURA and PURB may play regulatory roles in translation and splicing. Involvement in these processes is also supported by the high incidence of components of the translation and splicing apparatus in PURA and PURB protein-protein interactome (Figure 5I). It is also noticeable that both PURA and PURB datasets grouped together in the principal component analysis, with HIV-1 and mock treatment and not the protein identity as the main differential factor (Figure S8D). Moreover, 81% of the identified RNAs are targets for both proteins, displaying near identical binding patterns across their mRNA targets and a high degree of overlap of their binding sites (Figure 6A-F and S8F). These results reinforce the idea that PURA and PURB play cooperative or redundant roles in RNA metabolism.

**Figure 6.**
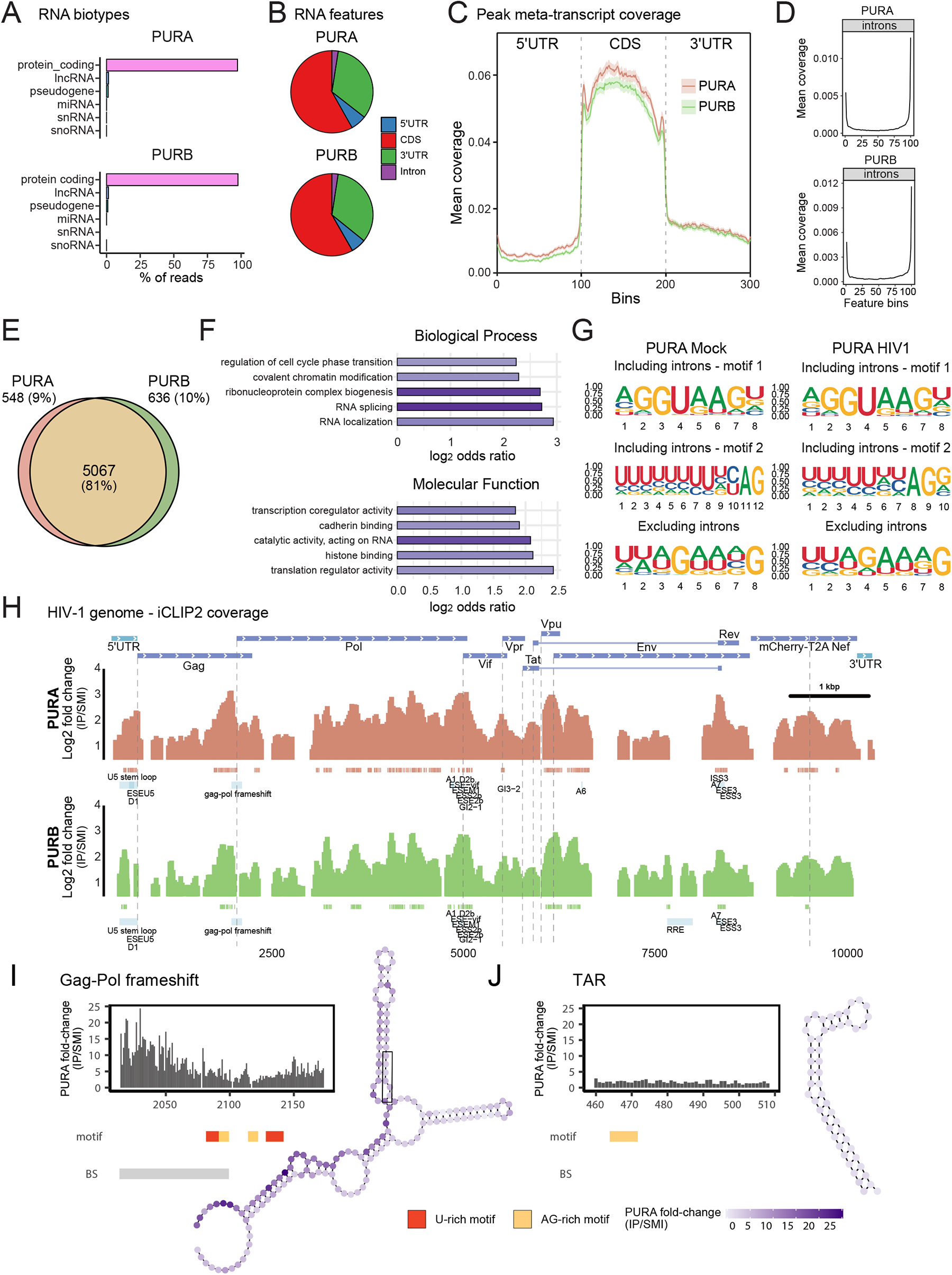
iCLIP2 analysis of PURA and PURB RNA targets. iCLIP2 was applied to Jurkat Flp-In T-REx expressing PURA-eGFP or PURB-eGFP and infected with HIV-1R-E-mCherry-Nef or mock-infected. A) Bar plots showing the biotypes of the RNAs identified by iCLIP2. B) Pie plots showing the distribution of binding sites within regions of the target RNAs. C) Density plot showing the distribution of binding sites across 5’ UTRs, CDSs and 3’ UTRs of target RNAs. D) As in (C) but for introns. E) Venn diagram comparing the RNA targets of PURA-eGFP and PURB-eGFP. F) GO terms enriched within the PURA-eGFP and PURB-eGFP RNA targets. G) Motifs enriched in RNA binding sites of PURA including or excluding intronic sequences. H) Binding profile of PURA-eGFP and PURB-eGFP on HIV-1 RNA. Only regions with >2-fold enrichment over size matched input (SMI) controls are shown. Significant binding sites (p<0.01) are indicated with pink (for PURA) or green (for PURB) boxes underneath the binding site density plot. I-J) Binding site density at the TAR (I) and Gag-Pol frameshift (J). The significant (p<0.01) binding sites are indicated with a grey box, while the AG- and U-rich motifs are indicated with yellow and red boxes, respectively. The secondary structure of these RNA elements are shown using SHAPE data, indicating the fold change in the IP over the SMI control at each nucleotide position.

To test if PURA and PURB recognise specific sequences, we examined the binding sites for enriched motifs. We found two prominent classes, a purine-rich motif that correlates well with the known binding preferences of PURA^49^, and a U-rich motif that was not previously described (Figure 6G and S8G). Detailed analysis of the occurrence of these motifs revealed that the U-rich motif precedes the AG-rich motif (Figure S8H). These motifs are highly similar to the canonical 5’ and 3’ splice sites^50^, which explains the distribution of PURA and PURB at the two ends of introns (Figure 6D). To rule out that these motifs are not an artifact derived from the proximity of PURA/B binding sites to splicing junctions, we searched again for motifs after removal of all intronic binding sites. In these conditions a single motif combining the U-rich sequence followed by the AG-rich sequence was observed (Figure 6G and S8H). Altogether, these data suggests that PURA and PURB interact with a bi-partite motif composed by a U-rich tract followed by an AG-rich sequence that is present at key regulatory elements within mRNAs.

### PURA and PURB bind to functionally important sequences within the HIV-1 gRNA

To determine if PURA and PURB interact with HIV-1 RNA, we searched for the binding sites mapping to the NL4.3 HIV-1 genome. 5-20% of the reads in HIV-1 infected cells mapped to the HIV-1 genome, which was substantially higher than those in controls (size matched inputs, SMI, Figure S8I). There was a high density of binding sites distributed across the gRNA with several high confidence binding sites located at important regulatory elements (Figure 6H). We observed a high density of reads at most canonical splice sites and splicing regulatory sequences in the gRNA (Figure S9), implying a potential involvement PURA and PURB in splicing regulation. Binding sites in splicing junctions contained, in most cases, the AG-rich or/and U-rich motifs (Figure S9). Moreover, a prominent PURA/B binding site was placed just before the slippery sequence that causes the Gag/Pol frameshift, which contained 2 repeats of the AG- and U-rich motifs that is compatible with the binding of more than one molecule of PURA/B (Figure 6I). However, we did not observe a substantial difference in Gag/Pol ratio in PURA and PURB KO cells, suggesting that these RBPs might not regulate frameshift efficiency (Figure S6G). While PURA was initially related to Tat-dependent regulation of HIV-1 transcription, we did not observe binding sites within the TAR structure that is the substrate for Tat (Figure 6J). PURA and PURB displayed nearly identical binding distribution across the gRNA, with a notable exception that is the binding of PURB to the RRE that was not as prominent for PURA (Figure 6H). Another binding site downstream the RRE and overlapping with several 3’ splice sites was detected for both PURA and PURB. Binding to RRE and its proximities is compatible with the efficient co-purification of these proteins in the Rev IP (Figure 3). Altogether, these results confirm that PURA and PURB interact with the HIV-1 gRNA at multiple key regulatory sequences, although the exact roles of these binding preferences in viral gene expression and RT deserve further characterisation.

## DISCUSSION

Many approaches have been previously undertaken to reveal the composition of viral particles^7^. However, the fact that virions assemble in the crowded cellular environment allows the passive incorporation of cellular proteins with no roles in the viral life cycle. Since viral particle proteomes are usually large and prone to false positives, the distinction between bystander and functionally relevant host proteins becomes a challenge. This is clearly shown in our study, as the HIV-1 particle proteome (ivRIC input) correlates well with protein abundance. By contrast, enrichment of the ivRNP by ivRIC allows the identification of proteins directly interacting with the viral RNA inside virions. Since the ivRNP components display a poor correlation with protein abundance, the incorporation of these proteins into virions is likely to occur through active mechanisms. The high performance of ivRIC is supported by the high number of previously established regulators of HIV-1 particle formation and infectivity that are top candidates in our datasets (e.g. PACSIN2^27^, UPF1^14^, EEF1A^15, 19^, MOV10^23, 24^, PDCD6IP [ALIX]^12^, APOBEC^13^ amongst others), as well as all proteins tested functionally here that showed effects in HIV-1 infection (i.e. MSN, FAM120A, PURA and PURB). Strikingly, we observed a strong overlap between the ivRNP and the Rev interactome, invoking an exciting link between the nuclear life of the HIV-1 gRNA and the formation of the viral particles. How these nuclear ivRBPs are captured and retained in the HIV-1 gRNP and whether they play a role prior to or after virus assembly requires a case-by-case study. In the present work we show that the ivRBPs MSN has no effect in virus infectivity but regulate viral gene expression and/or viral particle formation in the producer cell (Figure 5). However, other ivRBPs such as UPF1^14^, EEF1A^15, 19^, MOV10^23, 24^, APOBEC^13^, and, as shown here, PURA and PURB (Figure 5) aid or supress virus infectivity. In the future, ivRIC can be expanded to different producer cells, experimental conditions, and viruses to provide new insights into the cellular proteins that regulate the biology of viral particles. We applied here ivRIC to a virus with a polyadenylated genome; however, it can potentially be adapted to viruses lacking poly(A) tail by using specific antisense probe sets or total RNA purification methods^44^.

In the early 2000s, PURA was initially linked to Tat-dependent transcription using LTR-controlled plasmids^31^. However, these early publications were refuted by following studies as reviewed in detail elsewhere^51^. A major problem was the use of PURA constructs that, considering subsequent crystal structures, violated domain boundaries likely leading to unfolded and/or dysfunctional proteins. In addition, the studies were conducted using experimental approaches that do not allow separation of transcriptional and posttranscriptional processes^31, 51^. Here, we show that both PURA and PURB interact with cellular machineries involved in posttranscriptional regulation of gene expression and bind to key elements on the HIV-1 gRNA with central importance in posttranscriptional regulation, including the splice sites, the Gag/Pol frameshift site, start and stop codons, and the RRE. PURA and PURB have binding sites across the 5’ UTR with no overlap or proximity to the TAR structure that is bound by Tat. While results are not necessarily incompatible with the idea of a role for PURA in transcription, they suggest broader roles in posttranscriptional control.

An open question is how PURA and PURB regulate HIV-1 gene expression. Our data show several lines of evidence suggesting that PURA and PURB associate primarily with the unspliced gRNA: i) they are assembled with the gRNA into viral particles; ii) their binding sites span intron-exon junctions that are only found in unspliced HIV-1 RNAs; and iii) their binding density is similar in the Gag and Pol ORFs (present only in gRNA) to other parts of the genome (present also in spliced HIV RNAs). Since PURA and PURB bind to nearly all splice sites within the viral genome and interact with several splicing factors, it is plausible that these proteins regulate intron/exon recognition, probably inhibiting gRNA splicing. A role promoting the expression of unspliced gRNA, is compatible with the reduction of Gag and Gag-Pol levels observed in PURA KO cells. Moreover, PURA and PURB also bind to the Gag and Gag-Pol start and stop codons, the Gag/Pol frameshift site and several proteins involved in translation, suggesting an additional role in translation control. However, no differences in Gag/Pol ratio were detected, suggesting that the function of PURA/B might not be associated with the frameshift efficiency. The role of PURA and PURB in HIV-1 gene expression should be characterized with molecular detail in the future.

The function of PURA and PURB is not restricted to the producer cell but extends to the infectivity of the viral particles. The ivRBP EEF1A was proposed to modulate reverse transcription through its interaction with the RT^15^. PURA and PURB also interact with RT, the RNase H domain (p15) and IN. PURA and PURB could also contribute to RT directly through the interaction with the RT or indirectly through their activity on the gRNA. The fact that PURA is able to interact with both RNA and DNA^31^ is compatible with a regulatory role involving also the proviral DNA. IN has been proposed to play roles that span different stages of viral infection, particularly integration and Tat-dependent transcription^52^. Our results suggest that the regulatory roles of cellular proteins such as PURA and PURB may also span different stages of HIV-1 infection, affecting HIV-1 gene expression and viral particle infectivity.

## MATERIALS AND METHODS

### Contact for reagent and resource sharing

Further information and requests for resources and reagents should be directed to the Lead Contact, Alfredo Castello.

## VIRUS AND CELLS

### Cell culture

The following human cell lines are available in cell culture collections or commercially: T-lymphoblast SupT1 (ECAAC, #95013123), T-lymphocyte Jurkat Flp-In (Thermo Fisher Scientific, #R76207) and HeLa (ATCC, #CCL-2). HEK293T were kindly provided by Prof. Jan Rehwinkel (University of Oxford, UK). Jurkat Flp-In and tetracycline inducible Jurkat Flp-In T-REx expressing tagged RBPs were obtained as described in ‘Transfection and generation of inducible expression cell lines’. SupT1, Jurkat Flp-In and Jurkat Flp-In T-REx were cultured in RPMI-1640 with 1 mM sodium pyruvate and 10mM HEPES, while HEK293T were cultured in DMEM. Cell culture media was supplemented with 10% foetal bovine serum (FBS), 1x penicillin/streptomycin (Sigma Aldrich, #P4458), 1.25 µg/ml amphotericin B (Sigma Aldrich, #A2942) and the following specific antibiotics: 100 µg/ml Zeocin (Thermo Fisher Scientific, #R25001) for Jurkat Flp-In; 100 µg/ml Zeocin and 7.5 µg/ml Blasticidin S Hydrochloride (Cambridge Bioscience, #B001-100mg) for Jurkat Flp-In T-REx; 350 µg/ml Hygromycin B (Cambridge Bioscience, #H011-20ml) and 7.5 µg/ml Blasticidin for inducible Jurkat Flp-In T-REx eGFP, PURA-eGFP and PURB-eGFP. All cells were cultured in a humidified incubator at 37°C with 5% CO_2_.

### Viruses

Full length, infectious HIV-1_mCherry-Nef_ was obtained by transfection of pNL4.3-mCherry-T2A-Nef plasmid into HEK293T. For ivRIC, this primary stock was used to infect SupT1 cells so viral particles recovered for purification and UV crosslinking were produced in CD4+ T lymphocytic cells. The different pseudotyped HIV-1 were generated by co-transfection of HEK293T cells with pNL4-3.R-E-derived plasmids (Key Resources Table) and a plasmid encoding the vesicular stomatitis virus glycoprotein (pHEF-VSVG, NIH AIDS Reagent Program, #4693). Transfection and virus production methods are described below.

## MOLECULAR AND CELLULAR BIOLOGY AND VIROLOGY METHODS

### Plasmids and recombinant DNA procedures

To generate a stable mCherry-expressing, replication competent HIV-1 (named HIV-1_mCherry-Nef_), we followed a strategy analogous to Edmonds et al^53^. A synthetic DNA sequence containing mCherry followed by a T2A peptide coding sequence flanked by HIV-1 sequences was generated (GeneArt, Thermo Fisher Scientific). HIV-1 flanking sequences were derived from clone pNL4.3, starting at the unique BamHI site to the end of Env gene. A short DNA linker was introduced at the end of Env, including a consensus Kozak sequence before the mCherry start codon. At the end of the mCherry-T2A fusion, an unique XbaI site was introduced before continuing in frame with the start codon of Nef, finishing at the unique XhoI site within Nef. The resulting construct was cloned into pNL4.3 using the BamHI and XhoI unique restriction sites, generating pNL4.3-mCherry-T2A-Nef. The pcR-Blunt-NL4.3-mCherry-T2A-Nef plasmid, used for in vitro transcription of HIV-1 genomic RNA which was employed as standard in virus titration by RT-qPCR (see below), was generated by cloning the fragment between FspAI and PdiI from pNL4-3.mCherry-T2A-Nef into pcR-Blunt vector (Thermo Fisher Scientific, #K270020).

Rev-tagged plasmids pNL4.3.R-E-mCherry-T2A-Nef-Rev-Flag-3xMyc and pNL4.3.R-E-mCherry-T2A-Nef-Rev-Halo were made as follows. A synthetic DNA sequence containing a BamHI site, 173 bp of the NL4-3 Rev C-terminal domain, a flexible glycine-serine linker (TCGGCCGGAGGA), the relevant tag, a stop codon and a HpaI site was generated and inserted into pNL4.3.R-E-mCherry-T2A-Nef (described in ^54^) using BamHI and HpaI restriction sites. pNL4.3.R-E-mCherry-T2A-Nef-ΔRev plasmid was made by introducing 2 point mutations (T5974C and T6041A) in the Rev ORF of the pNL4.3.R-E-mCherry-T2A-Nef plasmid. Firstly, pNL4.3.R-E-mCherry-T2A-Nef was restricted with BamHI-HF and EcoRI enzymes to generate a 2.7kb template containing the Rev ORF for subsequent PCRs. To generate the T5974C point mutation, PCR was performed firstly with primers T5974C_Primer_A and T5974C_Primer_D and next with primers T5974C_Primer_B and T5974C_Primer_C. PCR mixtures were extracted from agarose gel, isolating 2 fragments encoding the point mutation. PCR was next carried out without primers or DNA to anneal extracted fragments. After 7 cycles, T5974C_Primer_A and T5974C_Primer_B were added and remaining cycles completed to produce a contiguous fragment harbouring the T5974C mutation. PCR mixtures were gel-purified and ligated into the restricted into pNL4.3.R-E-mCherry-T2A-Nef backbone. This process was then repeated with primers targeting the T6041A point mutation.

Plasmids encoding RBP-tagged proteins for the generation of inducible cell lines were cloned by 1) amplification of the protein with specific primers from SupT1 cDNA or available template plasmids, and 2) insertion into a pcDNA5/FRT/TO-eGFP-linker vector containing the eGFP tag before (N-terminus tagging) the multicloning site. eGFP was followed by the flexible linker GGSGGSGG.

Single-guide (sg)RNA expression plasmids for CRISPR/Cas9-mediated knock out were generated by inserting annealed oligos into the BLADE plasmid as described in ^55^. All plasmids were validated by sequencing.

### Virus production

To generate infectious HIV-1_mCherry-Nef_, we transfected the pNL4.3-mCherry-T2A-Nef plasmid into HEK293T using calcium phosphate as described in ^56^. Virus-containing supernatants were collected 48 hours post-transfection (hpt), cleared by centrifugation (3000g for 10 min, 4°C), filtrated with a 0.45µm PVDF Stericup-HV system (Merck, #S2HVU01RE), and viral particles were precipitated with PEG 6000^56^. This primary virus stock was titrated by infecting SupT1 cells by spinoculation (1200g for 1 h at 37°C in a swing out rotor), inactivating the virus 48 hours post-infection (hpi) in formaldehyde 4% for 1 h at 4°C, and counting mCherry-expressing cells in a flow cytometer (Beckman Coulter CytoFLEX LX or BD FACSCalibur, using BD FlowJo software for data analysis). To produce the virus in SupT1, we infected cells at MOI 0.1 by spinoculation and expanded the infected culture by replacing the growth medium every 48 h, keeping the concentration of living cells at 1×10^6^ cells/ml. Viruses in the supernatant were purified as above.

To obtain samples for ‘in virion RNA-interactome capture’ (ivRIC), infected SupT1 cells (or uninfected as control) were mixed with uninfected cells at a 1:4 ratio and a concentration of 1×10^6^ cells/ml. Medium was replaced by centrifugation at 400g for 5 min, and cells were incubated at 37°C and 5% CO_2_ in a T175 flask with 120 ml of medium. At 48 hpi, cells were collected by centrifugation (400g for 5 min) and supernatant was kept. Cell pellet was washed in PBS 1X and lysed in 5 ml of RIC lysis buffer 1X (20 mM Tris-HCl pH 7.5, 0.5 M LiCl, 1 mM EDTA, 0.1% IGEPAL, fresh 0.5% LiDS wt/vol and fresh 5 mM DTT) during 1 h at 4°C. Cell lysates were homogenized by pipetting and passing them through a 27G needle several times, and frozen at -80°C for whole cell proteome analysis. Supernatants were further cleared by centrifugation (3000g for 10 min at 4°C), followed by filtration with 0.45µm PVDF Stericup-HV filters. Viral particles were purified on a 10% sucrose cushion (50 mM TrisHCl pH 7.4; 100 mM NaCl; 0.5 mM EDTA; 10% sucrose) at a 4:1 vol:vol ratio (supernatant:sucrose) in 50 ml conical tubes (Sarstedt, #62.546.254) by centrifugation at 10000g for 4 h, 4°C according to the protocol described previously^57^. Virus-containing pellet was resuspended in PBS 1X containing 150 mM NaCl and 50 mM HEPES at a concentration ratio of 1:500 with respect to the initial volume, incubated overnight at 4°C, and finally aliquoted and stored at -80°C until use. The supernatant from uninfected cells was treated in an identical manner. Titration was performed by reverse-transcription and quantitative PCR (RT-qPCR) as described below. Alternatively, virus used for functional assays was obtained in SupT1 cells using a similar approach but concentrated with PEG 6000 and titrated by flow cytometry (see above).

Single-round, pseudotyped HIV-1 particles were produced by co-transfecting HEK293T cells with pNL4-3.R-E-derived plasmids together with pHEF-VSV-G using calcium phosphate. Virus was collected 48 hpt as before^56^ and titrated by flow cytometry.

### Virus titration by RT-qPCR

The number of viral particles employed in ivRIC was calculated as follows. 100 µl of purified HIV-1_Nef-mCherry_ (or supernatant from uninfected cells as control) was mixed with 100 µl of RIC lysis buffer 2X (40 mM Tris-HCl pH 7.5, 1 M LiCl, 2 mM EDTA, 0.2% IGEPAL, fresh 1% LiDS wt/vol and fresh 10 mM DTT) and incubated for 1 h at 4°C to inactivate the virus. Poly(A) RNA was purified using 100 µl of oligo(dT)_25_ beads (New England Biolabs, #S1419S) following the procedure described in the ‘ivRIC’ section but eluting in 100 µl of buffer and doing only one round of capture. Eluted RNA concentration was measured in a Qubit 2.0 Fluorometer using Qubit RNA High Sensitivity Assay Kit (Thermo Fisher Scientific, #Q32855), and the amount of HIV-1 RNA was assessed in a CFX96 Real-Time PCR Detection System (Bio-Rad) using Luna Universal One-step RT-qPCR kit (New England Biolabs, #E3005S) with primers specific to HIV-1 5’-UTR and the start of *gag* gene. 1:1, 1:10 and 1:100 dilutions were evaluated for each sample. HIV-1 RNA copy number was calculated in reference to a standard curve of purified HIV-1 gRNA, considering that each virion contains 2 copies of viral RNA. The standard curve was prepared as follows: 1) linearize pcR-Blunt-NL4.3-mCherry-T2A-Nef with MluI restriction endonuclease; 2) transcribe HIV-1 genomic RNA in vitro with HiScribe T7 ARCA mRNA kit (New England Biolabs, #E2065S); 3) add poly(A) tail with *E. coli* Poly(A) Polymerase (New England Biolabs, #M0276S); 4) purify RNA using oligo(dT)_25_ beads^58^; 5) estimate RNA concentration using Qubit RNA assay; 6) run 10-fold serial dilutions in the RT-qPCR in parallel to the samples under study; and 7) obtain curve Ct vs starting quantity, using the formula: Number of RNA copies = [1×10^-9^ g/ng * 6.022 * 10^23^ molecules/mol * amount of ssRNA in ng] / [(An x 329.2) + (Un x 306.2) + (Cn x 305.2) + (Gn x 345.2) + 159 g/mol]^59^. RT-qPCR data was analysed with CFX Manager Software v3.1 (Bio-Rad).

### *In virion* RNA interactome capture (ivRIC)

ivRIC was built on the previously established RNA-interactome capture protocol^58, 60, 61^ with some important modifications. 7×10^11^ purified HIV-1_mCherry-Nef_ particles per condition (or the equivalent volume of sucrose gradient purified supernatant from uninfected cells)^57^ were spread on a 6-well plate in 1 ml of total volume (topped up with 1x PBS) on ice and were irradiated with 150 mJ/cm^2^ of UV light at 254 nm. 333 µl of RIC lysis buffer 4X (80 mM Tris-HCl pH 7.5, 2 M LiCl, 4 mM EDTA, 0.4% IGEPAL, fresh 2% LiDS wt/vol and fresh 20 mM DTT) was then added to achieve 1X. Samples were recovered using a rubber cell scraper, transferred to a tube and homogenised by pipetting. Lysates were incubated for 1 h at 4°C to ensure the full inactivation of the virus, and then frozen down to -80°C for storage.

Samples were thawed at room temperature, and 10% of the sample was taken as input (i.e. proteome of viral particles). The rest of the sample was incubated under gentle rotation for 2 h at 4°C with 450 µl of oligo(dT)_25_ beads, previously equilibrated with RIC lysis buffer. Beads were collected in the magnet and the supernatant was stored at 4°C in a new tube to be used in a second round of oligo(dT)_25_ capture. Beads were washed once with 1.5 ml of 1X RIC lysis buffer for 5 min at 4°C, inverting the tubes ten times every minute. Beads were subsequently washed twice with 1.5 ml of cold buffer 1 (20 mM Tris-HCl pH 7.5, 500 mM LiCl, 0.1% LiDS wt/vol, 1 mM EDTA, 0.1% IGEPAL and 5 mM DTT), twice with 1.5 ml of cold buffer 2 (20 mM Tris-HCl pH 7.5, 500 mM LiCl, 1 mM EDTA, 0.01% IGEPAL and 5 mM DTT) and finally twice with 1.5 ml of room temperature buffer 3 (20 mM Tris-HCl pH 7.5, 200 mM LiCl, 1 mM EDTA and 5 mM DTT). Beads were resuspended in 200 µl of elution buffer (20 mM Tris-HCl pH 7.5 and 1 mM EDTA) and incubated for 3 min at 55°C with agitation (200 rpm). Beads were recycled in 0.1M NaOH for 5 min at 55°C with agitation (200rpm), equilibrated in 1X RIC lysis buffer, and re-used for one additional capture following the same protocol. Eluates of the first and second round of oligo(dT) capture were combined and stored at -80°C. Total eluted RNA was quantified using the Qubit RNA High Sensitivity Assay.

For quantitative proteomics, samples were treated with RNase A/T1 mix for 1.5 h at 37°C and then 15 min at 50°C as in^58^. Whole cell lysates were treated with 250U/ml of benzonase (Millipore, #70746-4) for 30 min. Protein concentration was measured using Pierce 660m Protein Assay Reagent and Ionic Detergent Compatibility Reagent (Thermo Fisher Scientific, #22660 and #22663 respectively) on a microplate reader (CLARIOstar Plus, BMG Labtech). Samples were further processed by single-pot, solid-phase-enhanced sample preparation (SP3) and peptides were analysed by LC-MS/MS as described in^61^.

For conventional protein analyses, ivRIC eluates were concentrated on an Amicon Ultra-0.5 centrifugal filter unit 3KDa cut-off (Millipore, #UFC500324) by centrifugation for 20 min at 18000g and 4°C. Filters were washed with 500 µl of 40 mM NaCl followed by centrifugation for 45 min at 18000g and 4°C. Proteins were recovered by placing the filter unit upside down in a new tube and spinning for 2 min at 1000g. Finally, RNA was digested with RNases T1 and A as above.

### Conventional protein analyses

Samples were mixed with NuPAGE LDS Sample Buffer 4X (Thermo Fisher Scientific, #NP0008), incubated for 10 min at 70°C, resolved by SDS-PAGE and analyzed by 1) Western Blot using specific antibodies (Key Resources Table), LI-COR Odyssey Fc imaging system for visualization and the Image Studio Software for quantification, or 2) silver staining using the SilverQuest kit (Invitrogen, #LC6070).

### HaLo labelling and single molecule RNA (sm)FISH

High Precision Coverslips (Marienfeld, #0107052) were washed once in 1 M HCl for 30 min on a rocking machine, twice in double distilled water for 10 min and once in ethanol 70% for 10 min. 150,000 HeLa cells were seeded on the prewashed coverslips in wells of a 6-well plate and incubated in DMEM with 10% FBS. Cells were infected 24 hours later using VSV-G pseudotyped single round virus (HIV-1_-R-E-Rev-HaLo_, HIV-1_-R-E-Δ Rev_ or HIV-1_-R-E-mCherry-Nef_). At 48 hpi, cells were washed with PBS and fixed with 4% paraformaldehyde for 10 minutes at room temperature. Cells were washed with PBS three times with gentle rocking and permeabilised with PBS + 0.1% Triton-X (PBSTX). Cells were washed three times with PBSTX for 5 minutes with gentle rocking. Janelia Fluor Halo-646 ligand (Promega, #GA1120) was diluted to 50 nM in PBSTX and dropped onto parafilm on a flat surface. Coverslips with attached cells were incubated face-down in PBSTX-dye mix for 1 h at room temperature. Coverslips were gently transferred to 6 well plates and washed 3 times with PBSTX before incubation for 5 min at 37°C with 2 μg/ml DAPI in PBSTX for 5 minutes. Finally, cells were washed twice with PBSTX, once with PBS for 5 min and once with milliQ H_2_O, followed by mounting on glass slides using Vectashield Antifade mounting medium (Vector Laboratories, #H-1000).

RNA FISH probes were designed using the LGC Biosearch Technologies’ Stellaris® RNA FISH Probe Designer. The first 2000 bp of the gag-pol ORF was targeted which is only present in gRNA. For smFISH, cells were seeded in coverslips and fixed and permeabilised as described above. Coverslips were washed at 37°C for 10 min with PBSTX, PBSTX with 1X saline-sodium citrate solution (SSC), PBSTX with 2X SSC and finally in pre-hybridisation buffer (2x SSC and 10% deionized formamide in DEPC water). Next, cells were incubated in a wet chamber for 16 h at 37°C with 125 nM HIV-1 gRNA-specific Stellaris probes (LGC Biosearch Technologies) in hybridization buffer (2x SSC, 10% deionized formamide and 10% dextran sulfate in DEPC water). Cells were subsequently washed twice with pre-hybridization buffer for 10 min at 37°C and incubated with DAPI and mounted as above. In both cases, images were acquired on an API DeltaVision Elite widefield fluorescence microscope using a 100X oil UPlanSApo objective (1.4 NA) and deconvolved with SoftWoRx v6.5.2 (GE Healthcare).

### Generation of knock-out SupT1 cells

To generate KO cells, we produced nanoblades loaded with Cas9 and specific sgRNAs targeting two regions for each gene as described before^47^. We then transduced 1×10^5^ SupT1 cells with 5 to 20 µl of nanoblades and 4 µg/ml polybrene in 200 µl of growth medium. 4 h later, 300 µl of fresh medium with 10% FBS was added to reach 500 µl. Cells were incubated for 48 h and then assessed for editing efficiency by genomic DNA extraction (Monarch Genomic DNA Purification Kit, New England Biolabs, #T3010S) followed by PCR, PCR cleanup (QIAquick PCR Purification Kit, Qiagen, #28104), sequencing and sequence trace decomposition using TIDE^62^. Single KO clones were obtained by serial dilution and validated by western blot and cytoplasmic mRNA quantification. For the latter, cytoplasmic RNA was extracted from 4×10^6^ cells by 1) lysing for 5 min on ice with 175 ul of chilled RLN buffer (50 mM Tris pH 8, 140 mM NaCl, 1.5 mM MgCl_2_, 0.5% IGEPAL); 2) spinning at 300g for 2 min at 4°C; and 3) purifying RNA from the supernatant using Monarch Total RNA Miniprep kit (New England Biolabs, #T2010). RNA quantification was performed by RT-qPCR using specific primers and Luna Universal One-step RT-qPCR kit.

### Analysis of cell viability and proliferation

To evaluate cell growth and viability, WT and KO cells were seeded at a concentration of 4×10^5^ cells/ml. 24, 48 and 72 h later, the number of cells and the percentage of living cells after trypan blue staining was estimated in a Countess II FL Automated Cell Counter (Thermo Fisher Scientific). At the same time points, cell viability was also assessed by adding CellTiter96 Aqueous One Solution Cell Proliferation Assay (Promega, #G3580) and measuring absorbance at 490 nm on a microplate reader.

### Analysis of HIV fitness

To assess if knocking out a protein affects virus gene expression and infectivity, we infected 2×10^6^ SupT1 cells (WT and KO) with infectious HIV-1_mCherry-Nef_ at 0.1 MOI by spinoculation, replaced growth medium at 2 hpi and incubated the cells for 48 h. Then, cells were collected by centrifugation at 400g for 5 min. 1) For viral gene expression assessment, cell pellet was resuspended in 2 ml of fresh medium (approximately 1×10^6^ cells/ml). 500 µl of infected cells were fixed in formaldehyde (4% final concentration), incubated 1 h at 4°C and analysed by flow cytometry. 2) The supernatant of these cells was collected and further cleared by centrifugation at 18000g for 10 min. Viral particles were precipitated with PEG 6000^56^ and titrated by RT-qPCR. 3) The remaining viral particle sample was normalised by RNA levels and used to infect 1.5×10^5^ SupT1 WT cells (second round) by spinoculation. After 48 h, cells were fixed and analysed by flow cytometer as before.

### Generation of Tet-on inducible Jurkat cells

To obtain the Jurkat Flp-In T-REx cell line, we first linearized pcDNA6/TR plasmid (Thermo Fisher Scientific, #V102520) with SapI restriction enzyme and transfected it into Jurkat Flp-In cells using a Lonza Amaxa Nucleofector II and the Cell Line Nucleofector Kit V (Lonza, #VCA-1003), according to the manufacturer’s recommendations for Jurkat E6-1 with the following modifications. Cells were seeded two days before nucleofection at a concentration of 2.5×10^5^ cells/ml in RPMI supplemented with 15% FBS without antibiotics. For nucleofection, 1×10^7^ cells (per reaction) were harvested by centrifugation (100g, 10 min), resuspended in 100 µl of Nucleofector Solution V plus Supplement and combined with 20 µg of plasmid. Nucleofection was performed using the program C-016. Immediately after, we added 500 µl of antibiotics- and FBS-free RPMI to each cuvette and incubated for 20 min at 37°C. Cells were transferred to a T25 flask containing 4.5 ml of RPMI with 20% FBS. Cuvettes were re-used for a second round of transfection with the same plasmid following the same procedure with these alterations: 1×10^7^ cells per reaction, 100 µl of Buffer 1SM (5 mM KCl; 15 mM MgCl_2_; 25 mM sodium succinate; 25 mM D-mannitol; 120 mM phosphate buffer Na_2_HPO_4_/NaH_2_PO_4_ pH 7.2) described in ^63^ and 40 µg of plasmid. Cells were mixed with those transfected in the first round and incubated for 48 h. Stably transfected single clones were selected by serial dilution in Zeocin plus Blasticidin containing media. Expression of Tet-R repressor was verified by Western blot. Repression and inducible expression were evaluated by nucleofection of pcDNA5/FRT/TO-eGFP plasmid, addition of 1 µg/ml doxycycline at 24 hpt and measurement of GFP signal at 48 hpt in a Countess II FL Automated Cell Counter.

To generate RBP-tagged inducible expression cell lines, we co-transfected Jurkat Flp-In T-REx cells with pOG44 (Thermo Fisher Scientific, #V600520) and the corresponding pcDNA5/FRT/TO plasmid (Key Resources Table) at a pOG44:pcDNA5/FRT/TO ratio of 8:2. At 48 hpt, zeocin was replaced by Hygromycin B for the selection of stable isogenic integrants. Protein expression was induced by adding 1 µg/ml doxycycline to the medium.

### Protein-Protein Interaction (Ppi) Analysis

#### The Rev interactome

VSV-G-pseudotyped HIV-1_-R-E-mCherry-Nef_ and HIV-1_-R-E-Rev-FLAG-Myc_ were produced as described above. 1.5×10^7^ SupT1 cells were seeded in 15 cm dishes and infected with either virus at 1 MOI. At 48 hpi, mCherry expression was assessed under a fluorescence microscope. Cells were pelleted by centrifugation at 400g for 5 min and resuspended in 1 ml ice-cold lysis buffer (50 mM Tris pH 7.5, 150 mM NaCl, 1% Triton-X, 0.5 mM EDTA, 25 U/ml of benzonase, 0.1 mM AEBSF) and lysed for 30 min on ice. Lysates were then briefly vortexed and centrifuged at 300g for 3 min. The supernatant was stored in 1.5 ml Eppendorf tubes. For pre-clearing, 100 µl of Pierce Control Agarose beads (Thermo Fisher Scientific, #26150) were washed three times with 1 ml of wash buffer (50 mM Tris pH 7.5, 150 mM NaCl, 0.2% IGEPAL, 0.5 mM EDTA) and incubated with the lysate for 30 min at 4°C with gentle rotation. Samples were then centrifuged at 2500g for 2 min and the supernatant was collected and transferred to a new tube. 40 µl of pre-equilibrated (in 1 ml of wash buffer for 1 h at 4°C with gentle rotation) anti-FLAG M2 magnetic beads (Merck, #M8823-1ML) were subsequently added to the sample and incubated for 1h. Beads were washed 6 times with 1 ml of wash buffer using a magnet For elution, beads were resuspended in a mixture of 20 µl of wash buffer and 10 µl 3X FLAG peptide (Merck, #F4799-4MG) and incubated for 1 h on ice The supernatant containing Rev-Flag-Myc was collected and stored. The elution process was repeated twice to maximise protein recovery.

#### PURA and PURB protein-protein interactions

Jurkat Flp-In T-REx PURA-eGFP and PURB-eGFP cells were induced with 1 µg/ml of doxycycline overnight. Cells were then infected with 1 MOI of VSV-G-pseudotyped HIV-1_-R-E-mCherry-Nef_ for 48 h and then lysed (10 mM Tris HCl pH 7.5, 150 mM NaCl, 0.5mM EDTA, 1% Triton X-100, 2mM MgCl2, 1mM DTT, 0.2 mM AEBSF serine protease inhibitor and 25 U benzonase). Cell lysates were cleared by centrifugation (17000 g, 10 min, 4°C) and protein concentration was measured using Pierce 660m Protein Assay Reagent and Ionic Detergent Compatibility Reagent. The lysate was pre-cleared with 40 ul of pre-equilibrated control agarose beads (Pierce Control Agarose resin, Thermo Fisher Scientific, #26150) by incubation under gentle rotation for 30 min at 4°C, followed by centrifugation at 2500 g for 2 min. Supernatants were transferred to a new tube and 2% of sample was taken as input. Samples were then incubated with 40 µl of pre-equilibrated GFP_Trap agarose bead slurry (ChromoTek, #gta) for 2 h at 4°C with gentle rotation. Beads were sedimented by centrifugation at 2500 g for 1 min at 4°C and washed six times with 500 µl of wash buffer (10 mM Tris HCl pH 7.5, 150 mM NaCl, 0.5 mM EDTA, 0.2% IGEPAL, 0.1 mM AEBSF serine protease inhibitor, 1mM DTT). Two additional wash steps were performed with 500 µl of wash buffer without IGEPAL. Proteins were eluted with 50 µl of 200 mM glycine pH 2.5 for 60 s followed by neutralisation with 5 µl of 1 M Tris base pH 10.4. Elution was repeated twice, and eluates were combined.

#### Determination of PURA and PURB binding sites on target RNAs by iCLIP2

The original iCLIP2 protocol^48^ was used with the following modifications. Jurkat Flp-In T-REx PURA-eGFP and PURB-eGFP cells were induced with 1 µg/ml of doxycycline overnight. Cells were then infected with 1 MOI of VSV-G-pseudotyped HIV-1_-R-E-mCherry-Nef_ for 48 h. Next, cells were washed twice in PBS 1X, resuspended in 3 ml of PBS 1X, crosslinked with two rounds of 0.15 J/cm^2^ UV light irradiation at 254 nm, washed with PBS 1X and lysed in 1 ml RIPA buffer (50 mM Tris pH 7.5, 150 mM NaCl, 1% Triton X-100, 0.1% SDS, 0.5% wt/vol Na deoxycholate and 0.2 mM AEBSF). Lysates were incubated for 30 min on ice and then stored at -80°C until use. Lysates were thawed on ice, sonicated with 3 cycles of 10 s at 4°C (with 15 s pause between pulses) using a Digenonde bioruptor at level M, homogenized by passing through a 27G needle several times, and finally cleared by centrifugation (17000 g for 10 min at 4°C). eGFP signal was measured on a microplate reader to normalize the amount of lysate for each sample.

4 U TurboDNase (Thermo Fisher Scientific, #AM2238) and 5 U RNse I (Thermo Fisher Scientific, #AM2294) were added, mixed with a vortex and incubated for 3 min at 37°C at 1100 rpm. 200 U RiboLock RNase Inhibitor (Thermo Fisher Scientific, # EO0381) was then added, followed by incubation for 3 min on ice. Lysates were pre-cleared with 25 ul of pre-equilibrated control agarose beads for 30 min at 4°C with gentle rotation followed by centrifugation at 2500 g for 2 min. Supernatants were transferred to a new tube and 2 x 10 µl of sample was taken for size-matched input (SMI) processing. Lysates were incubated with 25 µl of pre-equilibrated GFP_Trap agarose bead slurry for 2 h at 4°C with rotation. Beads were washed twice with 900 µl of cold high-salt wash buffer (50 mM Tris-HCl pH 7.4, 1 M NaCl, 1mM EDTA, 1% IGEPAL, 0.1% SDS, 0.5% sodium deoxycholate and 0.2 mM AEBSF), twice with 900 µl of cold medium-salt wash (20 mM Tris HCl pH 7.4, 250 mM NaCl, 0.05% IGEPAL, 1 mM MgCl_2_ and 0.2 mM AEBSF), and twice with 900 µl of cold PNK wash buffer (20 mM Tris HCl pH 7.4, 10 mM MgCl_2_ and 0.2% Tween-20). RNA 3’- end was dephosphorylated at 37°C for 40 min at 1100 rpm in PNK buffer (70 mM Tris HCl pH 6.5, 10 mM MgCl_2_ and 1 mM DTT) with 5 U PNK (New England Biolabs, #M0201L), 0.25 U FastAP alkaline phosphatase (Thermo Fisher Scientific, #EF0654), 0.5 U TurboDNase and 20 U RNasin (Promega, #N2111). Beads were washed once with 500 µl cold PNK wash buffer, twice with the same volume of cold high-salt wash buffer and twice with cold PNK wash buffer. 125 nM of L3-IR-App adapter^64^ was ligated using 30 U T4 RNA ligase I High Concentration (New England Biolabs, #M0437M), 20 U RNasin, 4 U PNK, 22.5% PEG8000 and 5% DMSO in ligation buffer (50 mM Tris HCl pH 7.8, 10 mM MgCl_2_ and 1 mM DTT) at 16°C overnight with shaking at 1100 rpm in the dark. Beads were washed once with 500 µl PNK wash buffer, twice with the same volume of high-salt wash buffer and twice with PNK wash buffer. IP samples and inputs were denatured in 1X

NuPage LDS Sample Buffer (Thermo Fisher Scientific, #NP0007) with 100 mM DTT for 5 min at 70°C, spun down for 2 min at 2500 g and separated on a NuPAGE 4-12% Bis-Tris gel (Thermo Fisher Scientific, #NP0321BOX) for 65 min at 180 V. Protein-RNA complexes were transferred onto an iBLOT2 nitrocellulose membrane (Thermo Fisher Scientific, #IB23001) for 2 h at 30 V and visualized on a LI-COR Odyssey Fc imaging system. The region corresponding to the RBP-eGFP band and above was cut (for both IP and SMI samples) and digested using ∼350 µg Proteinase K (Roche, #3115828001) in 180 µl PK-SDS solution (10 mM Tris HCl pH 7.4, 100 mM NaCl, 1 mM EDTA and 0.2% SDS) for 60 min at 50°C with shaking (1100 rpm). RNA was purified by adding 1X volume of Phenol:Chloroform:Isoamyl Alcohol pH 6.6-6.9 (Sigma-Aldrich, #P3803), incubating for 10 min at 37°C with shaking (1100 rpm) and phase separation in MaxTract tubes by centrifugation at 16000 g for 5 min. RNA was cleaned using Zymo RNA Clean & Concentrator-5 (ZYMO Research, #R1013). For SMI library preparation, SMI control samples were treated first with 5 U PNK, 0.5 U FastAP and 20 U RNAsin in PNK buffer pH 6.5 for 40 min at 37°C and 1100 rpm. RNA was cleaned up with Dynabeads MyOne Silane (Thermo Fisher Scientific, #37002D). L3-IR-App adapter ligation was performed with 45 U T4 RNA ligase I High Concentration in 1X T4 RNA Ligase Reaction Buffer with 2% DMSO, 27% PEG8000, and 133 nM L3-IR-adapter for 75 min at room temperature, followed by MyOne bead purification. SMI was treated with 25 U 5 ′ Deadenylase (New England Biolabs, #M0331S) and 15 U RecJf endonuclease (New England Biolabs, #M0264S) in 1X New England Biolabs buffer 2 with 20 U RNAsin, 20% PEG8000 for 1 h at 30°C and then 30 min at 37°C at 1100 rpm, followed by a MyONE clean-up. RNA from IP and SMI samples were reverse transcribed using Superscript IV reverse transcriptase (Thermo Fisher Scientific, #18090010) and hydrolyzed by adding 1.25 µl of 1 M NaOH for 15 min at 85°C before neutralization with 1.25 μl of 1 M HCl. cDNA was purified using MyOne silane beads. L#clip2.0 adapters with barcodes for multiplexing^48^ were ligated to cDNA by mixing first 2 μl of 10 μM adapter with 5 μl of cDNA and 1 μl of DMSO and incubating at 75°C for 2 min before placing on ice. Then, ligation mix (45 U T4 RNA ligase I High Concentration in 1X RNA ligase buffer with 1 mM ATP and 22.5% PEG8000) was added to the cDNA-bead solution and incubated overnight at 20°C with shaking 1100 rpm. cDNA was cleaned up with MyONE beads before PCR amplification. Pre-amplification was performed using 2X Phusion HF PCR Master mix (New England Biolabs, #M0531L) with P5Solexa_s and P3Solexa_s primers for six cycles, followed by ProNex (Promega, #NG2001) size-selective purification. Optimal qPCR cycles were determined on a CFX96 Touch Real-Time PCR Detection System (Bio-Rad) using EvaGreen (Biotium, #31000), 2X Phusion HF PCR Master mix and P5/P3 Solexa primers. Final PCR products were purified using two consecutive rounds of ProNex Size selection. Libraries were quantified by qPCR using the KAPA Library Quantification DNA standards (Roche, #07960387001) and High Sensitivity DNA kit (Agilent, #5067-4626) in an Agilent 2100 Bioanalyzer instrument. Each group of samples was pooled equimolarly and then mixed at the following proportions: 50% IP library pool, 37.5% SMI library pool, and 12.5% negative control eGFP. Sequencing was performed on a NextSeq 550 sequencer with a 75 cycle High-output kit v2.5 (Illumina, #20024906).

## PROTEOMIC AND TRANSCRIPTOMIC ANALYSES

### Sample preparation for mass spectrometry

ivRIC eluates were processed by single-pot, solid-phase-enhanced sample preparation (SP3) and peptides were analysed by LC-MS/MS as described in detail in ^61^. ivRIC samples and PURA-eGFP and PURB-eGFP IP were prepared for proteomics using SP3 as previously described^61^. Eluates of Rev IP were processed using Filter-Aided Sample Preparation (FASP) protocol^65^ with custom adjustments. IP eluates were denatured in 200 µl of Urea buffer (8 M Urea, 100 mM ammonium bicarbonate-AmBic-in water) by pipetting. Denatured samples were subjected to reduction and alkylation in solution by addition of 10 mM TCEP and 50 mM CAA, and incubated in dark for 30 min. VIVACON500 30 kDa filters were washed with 200 µl of 0.1% TFA, 50% ACN, and centrifuged at 14,000 g for 15 min. Alkylated samples were loaded onto 30 kDa filters and centrifuged at 14,000 g for 15 min. Detergent contaminants were removed by washing 3 times as follows: 200 µl of urea buffer was added to the filter unit and mixed with gentle pipetting; washing solution was passed through the filter by centrifugation at 14,000 g for 15min and discarded. Filters with the detergent-free samples were washed twice with 200 µl of 1 M Urea in 25 mM AmBic, and centrifugated at 14,000 g for 5 min. On-filter digestion was then performed by addition of 200 µl trypsin endoproteinase mixture (250 ng worth of trypsin, 1 M urea, 25 mM AmBic, 1 mM CaCl_2_), and incubated at 37°C overnight. Peptides were collected by filter centrifugation at 14,000 g for 15 min. The filters were washed first with 150 µl 0.1% Trifluoroacetic acid (TFA), and then 150 µl 50% acetonitrile (ACN) in 0.1% TFA, with a centrifugation at 14,000 g for 15min after each addition. Flowthrough fractions in these two washes were combined with the previous eluate to maximise peptide recovery. Combined flow-through were dried using SpeedVac and peptides were then reconstituted with MS loading buffer (5% DMSO, 5% FA in water) for LC-MS/MS analysis.

### LC-MS/MS analysis

Tryptic peptides were analysed using Ultimate 3000 ultra-HPLC system (Thermo Fisher Scientific) connected to a Q-Exactive mass spectrometer (Thermo Fischer Scientific) through an EASY-Spray nano-electrospray ion source (Thermo Fischer Scientific). The peptides were first trapped on C18 PepMap100 pre-column (300 µm inner diameter x 5 mm, 100 Å, Thermo Fisher Scientific) in solvent A (0.1% FA in water), and were then separated on the analytical column (75 µm inner diameter x 50 cm packed with ReproSil-Pur 120 C18-AQ, 1.9 µm, 120 Å, Dr. Maisch GmbH) using 30 min (ivRIC) or 60 min (PPI & SupT1 whole cell proteome) 15-35% (vol/vol) acetonitrile linear gradient at a flow rate of 200 nl/min. Full-scan mass spectra were acquired in the Orbitrap (scan range 350-1500 m/z, resolution 70000, AGC target 3 x 106, maximum injection time 50 ms) in a data dependent mode. Subsequently, the 10 most intense peaks were selected for higher energy collisional dissociation fragmentation at 30% of normalised collision energy. Higher-energy collisional dissociation fragmentation spectra were also acquired in the Orbitrap (resolution 17500, AGC target 5 x 104, maximum injection time 120 ms) with first fixed mass at 180 m/z.

Protein identification and quantification were performed using Andromeda search engine implemented in MaxQuant (1.6.3.4). Mass spectra were searched against human proteome reference (Uniprot_id: UP000005640, downloaded Nov 2016) and a custom HIV-1 (NL4-3) proteome including all known viral proteins. Flag-Myc-tagged Rev and eGFP were also included in the searches of the Rev and PURA/B interactomes, respectively. Search parameters included: full tryptic specificity with maximal two missed cleavage sites, carbamidomethyl (C) set as fixed modification, acetylation (protein N-term) and oxidation (M) set as variable modifications. False discovery rate (FDR) cut-off for peptide identification was set to 1%. For ivRIC and SupT1 whole cell proteome iBAQ and LFQ options were toggled ON. All other settings were set to default.

## QUANTIFICATION AND STATISTICAL ANALYSES

### Quantitative analysis of proteomics data

The proteinGroup files of MaxQuant search results were imported in RStudio (R Project) for further processing. Protein intensities were log 2 transformed. In each dataset proteins with less than 2 valid intensity measurements across experimental conditions were removed prior to downstream analysis. Batch effects in each comparison were assessed using principal component analysis (PCA).

Normalisation was performed to ivRIC inputs ad PPI analyses using variance stabilisation normalisation (vsn) method^66^. Missing values were imputed with deterministic minimum method (R package version 2.0. https://CRAN.R-project.org/package=imputeLCMD) using 1% quantile of global intensities. All missing values were imputed in ivRIC and Rev interactome data. For PURA/PURB interactome data, only proteins with all values missing in one condition were imputed as described in^67^. Linear modelling and Bayesian-model-based moderated T-test was performed using the R-package *limma*^68^. Batch effects in ivRIC eluates and Rev and PURA/B interactomes were accounted for by incorporation in modelling using “block” argument provided in *limma*. P values obtained in the moderated T-test were adjusted to account for multiple testing using Benjamini-Hochberg method.

### iCLIP2 data processing

The raw FASTQ files were demultiplexed according to the sample barcode using Je Suite (Girardot et al., 2016) and adapter trimmed with Cutadapt (Martin, 2011). Trimmed reads were aligned to a concatenated human (GRCh38, ENSEMBL Release 104) and HIV-1 NL4.3 genome using STAR with end-to-end alignment mode^69^. Only uniquely aligned reads were considered for the downstream analysis. PCR duplicated reads were collapsed using unique molecular identifiers (UMIs) attached to the read header with the Je Suite. The GRCh38 and HIV-1 genomic annotations were pre-processed to generate sliding windows (50nt window, 20nt step size) using HTSeq-clip^70^. Crosslink truncation sites (position -1 relative to the 5’ end of the read start) were extracted using BEDTools^71^ and quantified against the sliding windows using HTSeq-clip. For peak calling, a R/Bioconductor package DEW-Seq was used to identify significantly enriched sliding windows in PURA/PURB immunoprecipitated samples over the corresponding size-matched input control samples (log2FoldChange > 2 and p.adj < 0.01)^70^. The Independent Hypothesis Weighting (IHW) method was used for the multiple hypothesis correction^72^ To remove background signal resulting from non-specific binding of RNA to GFP, significantly enriched sliding windows (log2FoldChange > 2 and p.adj < 0.01) from GFP-immunoprecipitated control samples were removed. Overlapping significant sliding windows were merged to binding regions, and these sites were curated to 8nt long binding sites based on peak width and maxima. Binding sites were queried against the genome annotation (ENSEMBL release 104) using the GenomicRanges R package to extract overlaps with genes and transcript features^73^. The overlap information was used to construct the meta-transcript coverage of binding sites and the heatmap distribution of PURA/PURB binding in the proximity of start and stop codons.

Given the repetitive motifs present in 5’ and 3’ of HIV-1 genome, we used a custom transcriptome to identify binding motifs at the 5’ and 3’ untranslated regions (UTRs). In brief, we superimposed the LTR-1 and LTR-2 sequences flanked by 100bp of protein-coding sequences from HIV-1 open reading frames at both 5’ and 3’ ends. HIV-1 reads were initially filtered using BBMap with kmer = 25 mode (https://sourceforge.net/projects/bbmap/), then mapped to the custom transcriptome using STAR. Peak-calling was performed as described above using DEW-Seq except for using smaller sliding windows (10nt window, 2nt step size) and a more lenient threshold (log2FoldChange > 1.5 and p.adj < 0.01).

For the comparison of different iCLIP libraries, a principal component analysis was initially performed. First, the libraries were size-corrected and transformed using the variance stabilisation method implemented in DESeq2. Then, the 1,000 sliding windows showing the largest variance were used to perform the dimensionality reduction using the prcomp function implemented in the base R. To assess the spatial correlation of PURA and PURB binding sites, the relative distance metric was calculated using BEDTools reldist function. The same density of PURA and PURB binding sites was randomly simulated across the genome to compare with the observed nearest neighbour distance.

### Motif analysis in PURA and PURB binding sites

Sequences for motif enrichment analysis were defined for each binding site as a 70-nucleotide region, centred on the peak in BigWig signal. The sequence was extracted directly from UCSC genome hg38 when intron sequence was included in the analysis. When introns were excluded, sequence was extracted from the CDS sequence of the longest isoform of the gene in which the binding site occurred. For each binding site, a gene and gene region matched background sequence were defined to allow for differential enrichment. Enrichment analysis was performed using HOMER. Motifs were processed and plotted using the R packages universalmotif and ggseqlogo.

### Mapping and comparing gene IDs

For all datasets analyses, including mass spectrometry data, genes were mapped to Hugo gene nomenclature committee IDs (HGNC) using R package BiomaRt. Any genes that could not be mapped were manually curated by manual addition or were removed if it was a pseudogene or its HGNC entry was missing. For HIV-1 analysis, the HIV-1 NCBI database was downloaded from the NCBI web server and parsed to gene IDs following the same strategy. Upset plots and Euler diagrams were generated using the R packages upsetR and venneuler respectively.

### GO and STRING network analyses

General GO terms were extracted using the tool EnrichR^74^

(https://maayanlab.cloud/Enrichr/) and summarized using REVIGO^75^.

To generate virus GO term plots, the R package AnnotationDbi was queried with HGNC IDs and resulting GO terms filtered by the following GO categories: ‘viral’, ‘immune’, ‘infection’, ‘pathogen’, ‘immune cell’ and ‘immune molecule’.

Network analyses were performed using the Cytoscape 3.9.1 platform ^76^ with the following add-ons: stringApp ^77^ for STRING protein network analysis and GO enrichment; clusterMaker2 ^78^ for clustering data using MCODE algorithm ^79^ and DyNet^80^ for comparison of two networks.

### Classification of RNA-binding proteins

We classified proteins as RBPs if they were identified in at least 3 independent RNA interactome studies based on the EMBL RBPbase (https://rbpbase.shiny.embl.de/) resource.

## KEY RESOURCES TABLE

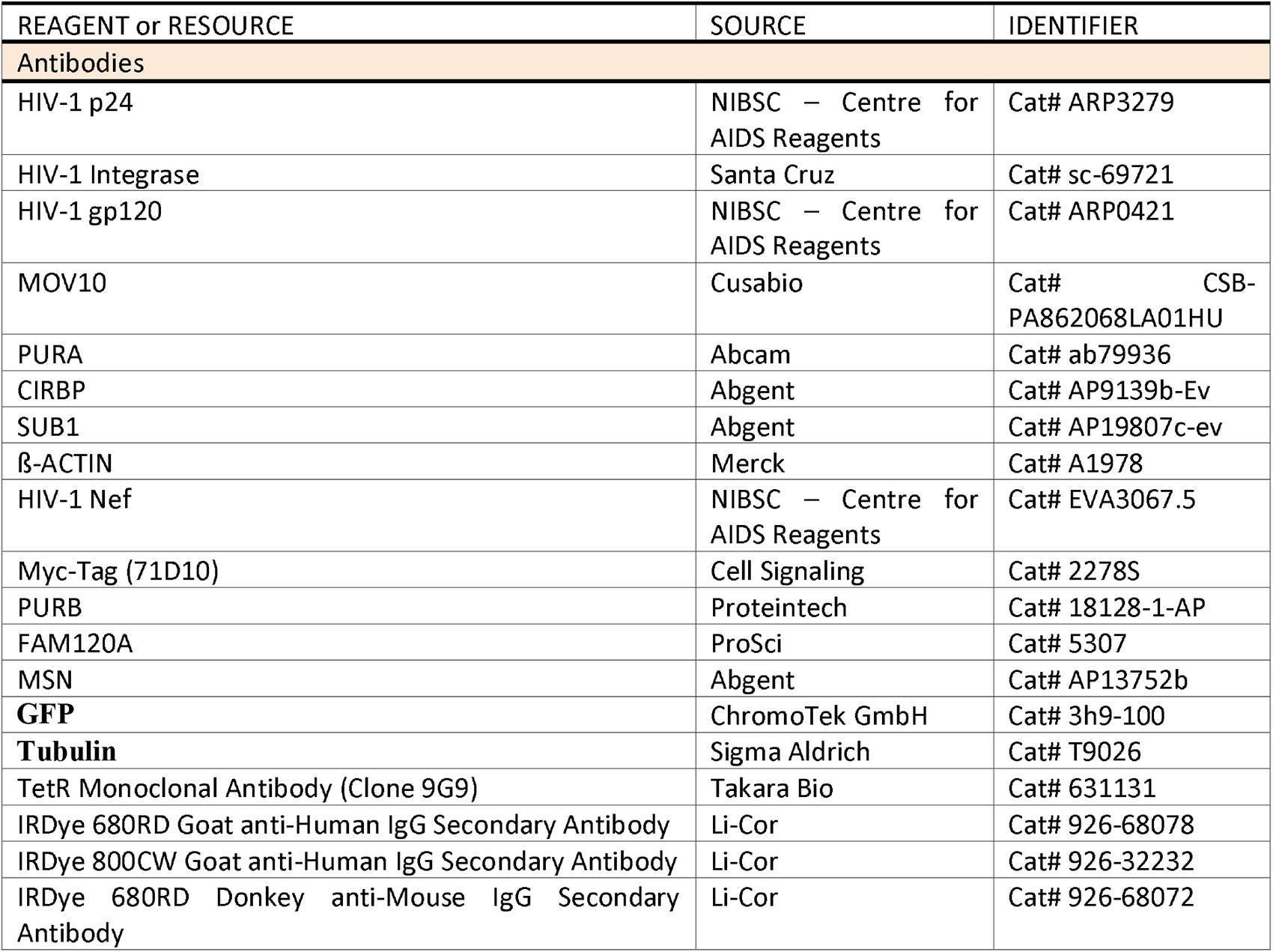
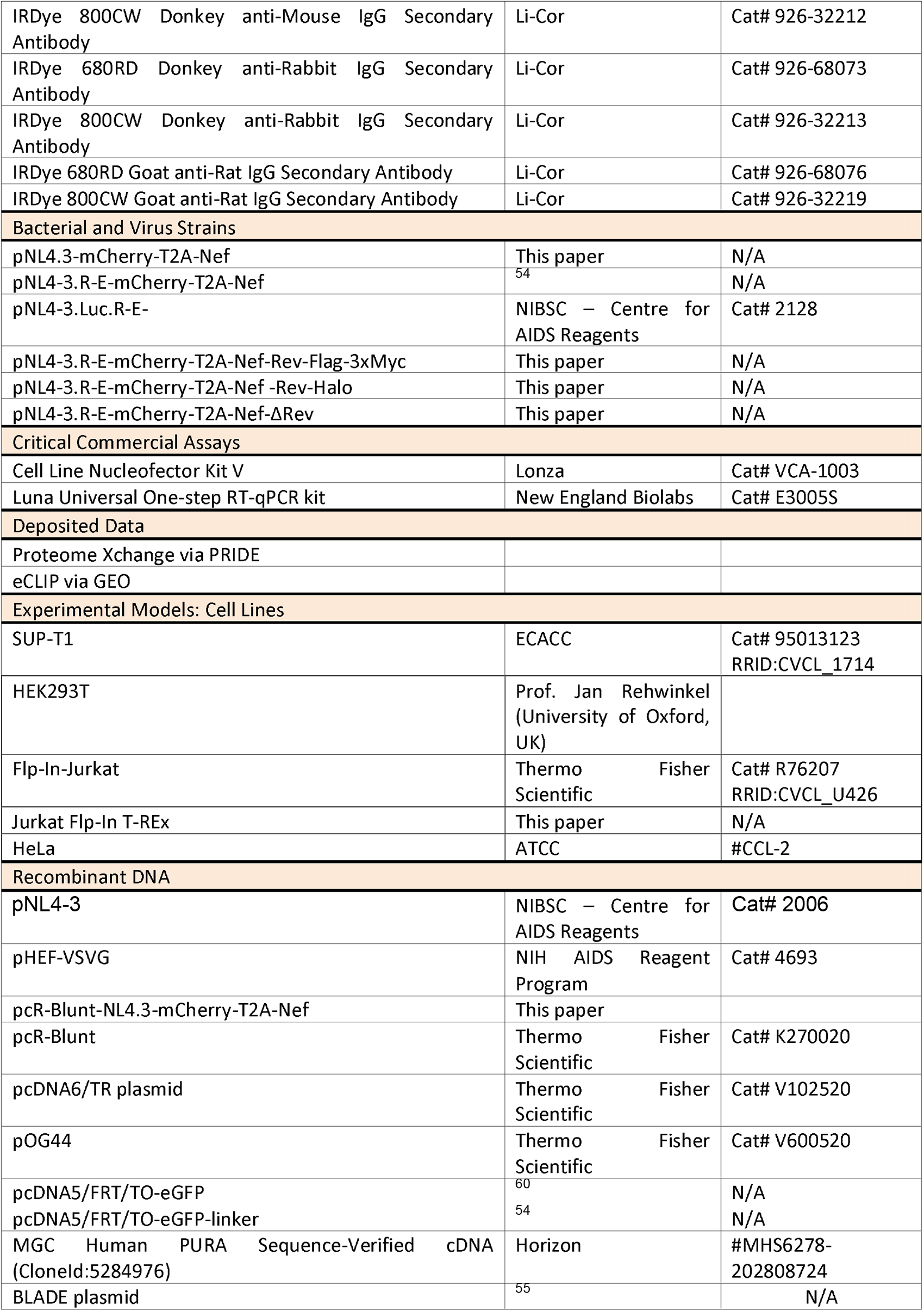

## ACKNOWLEDGEMENTS

AC is funded by the European Research Council (ERC) Consolidator Grant ‘vRNP-capture’ N# 101001634, the Career Development Award #MR/L019434/1, and the John Fell Funds from the University of Oxford. M.G.M is funded by the European Union’s Horizon 2020 research and innovation programme under the Marie-Sklodowska-Curie grant agreement N# 700184. R.T. is funded by a BBSRC DTP scholarship DD01.20. L.I. is funded by the BBSRC DTP scholarship number BB/M011224/1. C.E.L. is funded by the Department of Biochemistry Graduate Scholarship and the Hertford College Graduate Studies Scholarship of the University of Oxford. T.J.M.S is funded by Fondation pour la Recherche Medicale (FRM – FDT202001010798). W.K. is funded by the European Union’s Horizon 2020 research and innovation programme under Marie-Sklodowska-Curie n# 842067. V.R. is funded by the European Union’s Horizon 2020 research and innovation programme under Marie-Sklodowska-Curie n#892756. E.P.R is funded by Agence Nationale des Recherches sur le SIDA et les Hépatites Virales (ANRS – ECTZ3306), Fondation FINOVI and the European Research Council (ERC-StG-LS6-805500) under the European Union’s Horizon 2020 research and innovation programs as well as the ATIP-Avenir program. I.D. is funded by the Wellcome Trust Investigator Award 209412/Z/17/Z.

## AUTHOR CONTRIBUTIONS

Conception/design, M.G-M., R.T., A.C.; Data acquisition, M.G-M., R.T., C.E.L., H.C., M.N., T.J.M.S., K.D., R.P., N.J., V.R., S.H.; Data analysis, M.G-M., R.T., C.E.L., H.C., A.I.J., W.K., J.L., S.H., S. M., A.C.; Data interpretation, M.G-M., R.T., M.N., S.M., A.C.; Writing, original draft, A.C., M-G.M.; Writing, editing, all authors; Funding acquisition, M.G-M., I.D., S.M., A.C.; Resources, T.J.M.S., J.L., E.P.R., I.D., S.M.; Supervision, M.G-M., M.N., E.P.R., I.D., S.M., A.C.

