## Supplementary material for "Incorporation of genome-bound cellular proteins into HIV-1 particles regulates viral infection": Figure S1

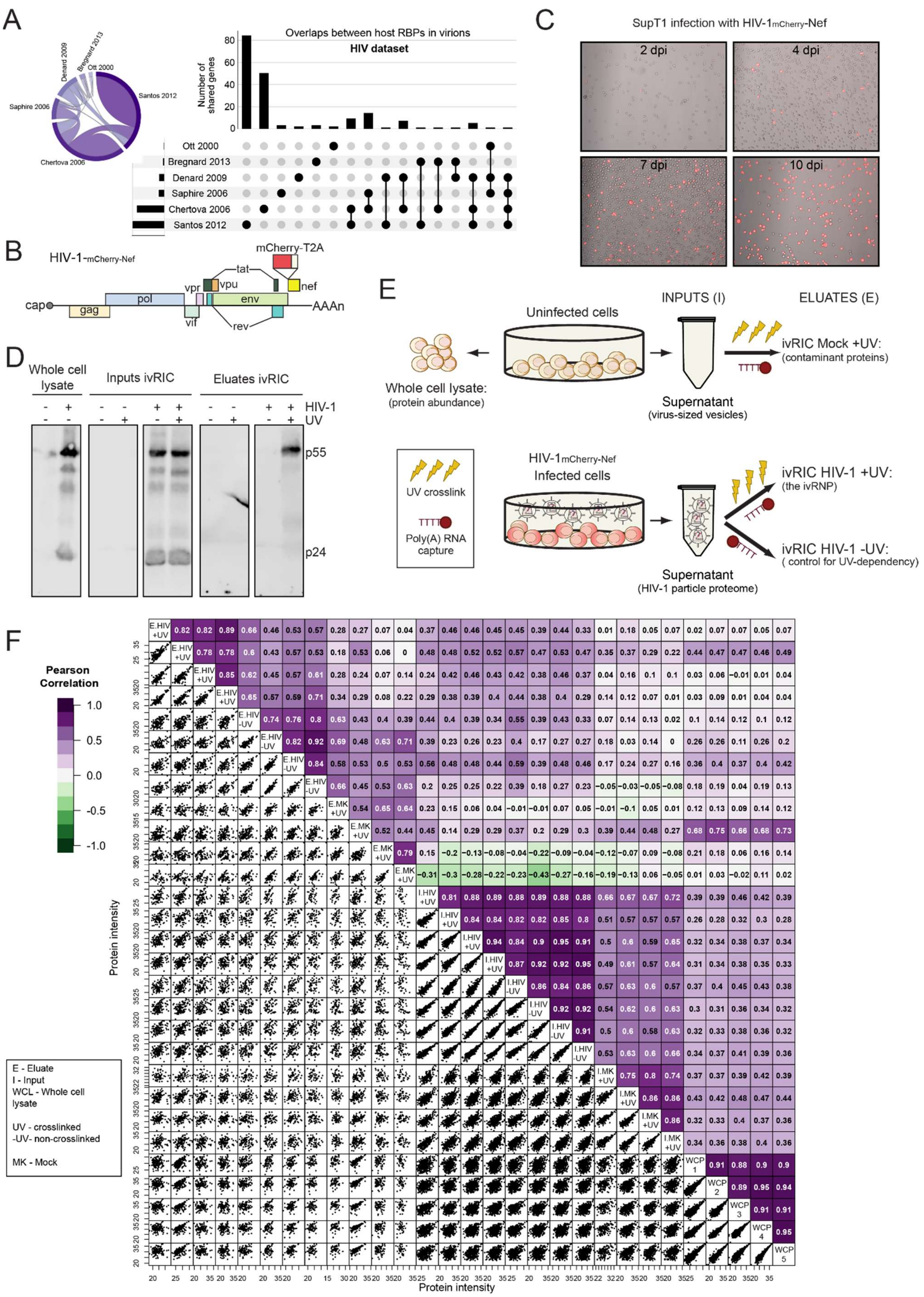


**Figure S1. Proteomic analysis of the ivRNP.** A) Comparison of previous HIV-1 particle proteomes. B) Schematic representation of the HIV-1$_{\text{mCherry-Nef}}$ chimeric virus. C) SupT1 cells were infected at 0.1 MOI with HIV-1$_{\text{mCherry-Nef}}$. The presence of mCherry-expressing cells was checked by fluorescent microscopy at different days post infection (dpi). D) Western blotting against CA/p24 in whole cell, input and eluate samples of an ivRIC experiment in HEK293T cells transfected with the plasmid encoding $HIV\text{-}1_{mCherry\text{-}Nef}$. E) Schematic representation of the ivRIC experimental design used for proteomics. F) Scatter plots showing the protein intensity and the Pearson correlation between different samples and replicates of the ivRIC experiment.
