## Supplementary material for "Incorporation of genome-bound cellular proteins into HIV-1 particles regulates viral infection": Figure S2

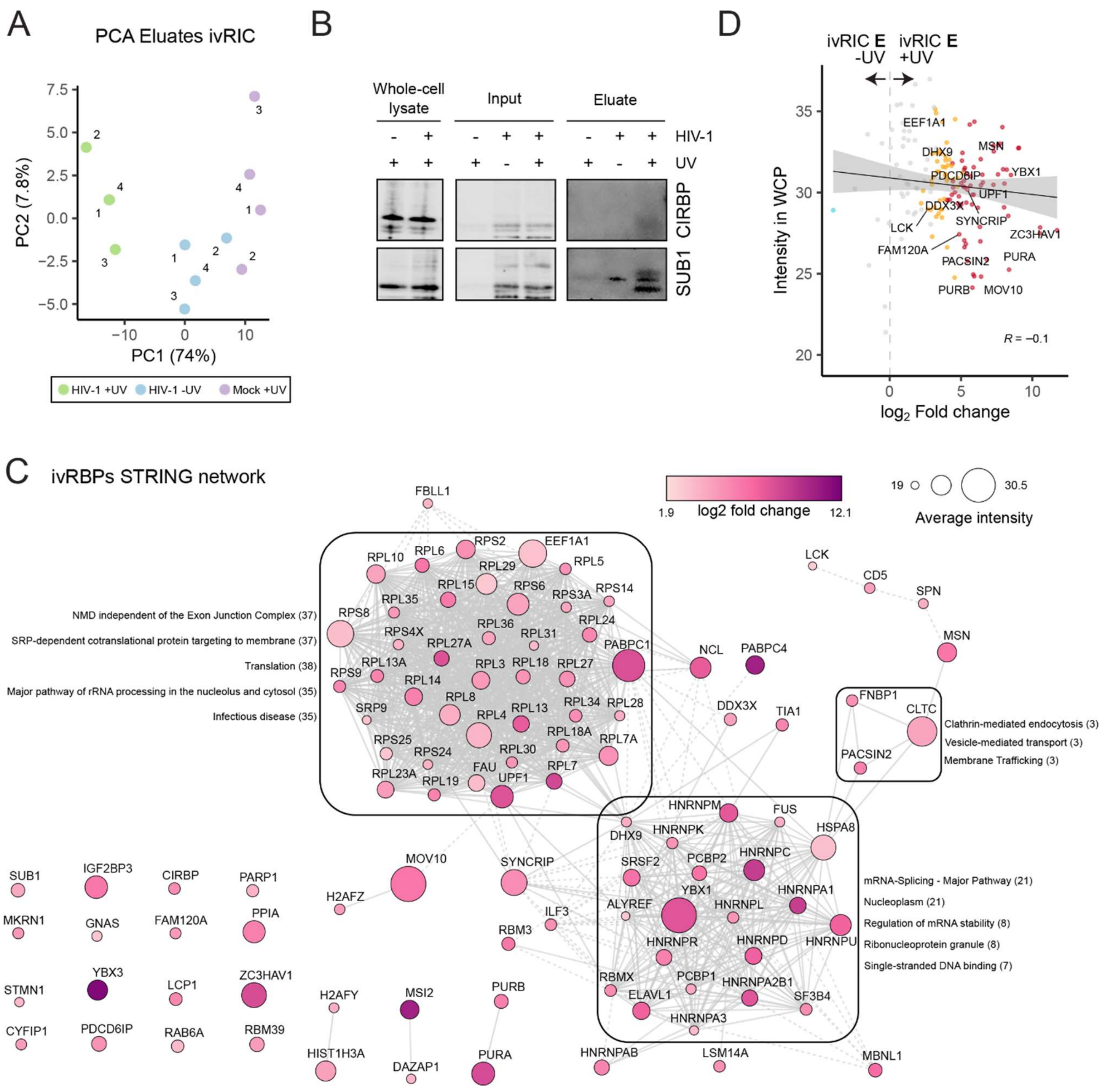


**Figure S2. Analysis of the ivRIC proteomic experiment in HIV-1-infected CD4+ T lymphocytic cells.** A) Principal component analysis (PCA) of the ivRIC eluates. B) Western blotting analysis of two discovered cellular ivRBPs in whole cell lysates, inputs and eluates of ivRIC. C) STRING network analysis of the ivRBPs with Cytoscape. Physical interactions are represented by solid lines and functional interactions by dashed lines. GO enriched terms are shown for each cluster. D) Scatter plot showing the relation between protein intensity in the whole cell proteome (WCP) and the fold change in UV irradiated versus non-irradiated HIV-1-infected samples of the ivRIC experiment.
