## Supplementary material for "Incorporation of genome-bound cellular proteins into HIV-1 particles regulates viral infection": Figure S3

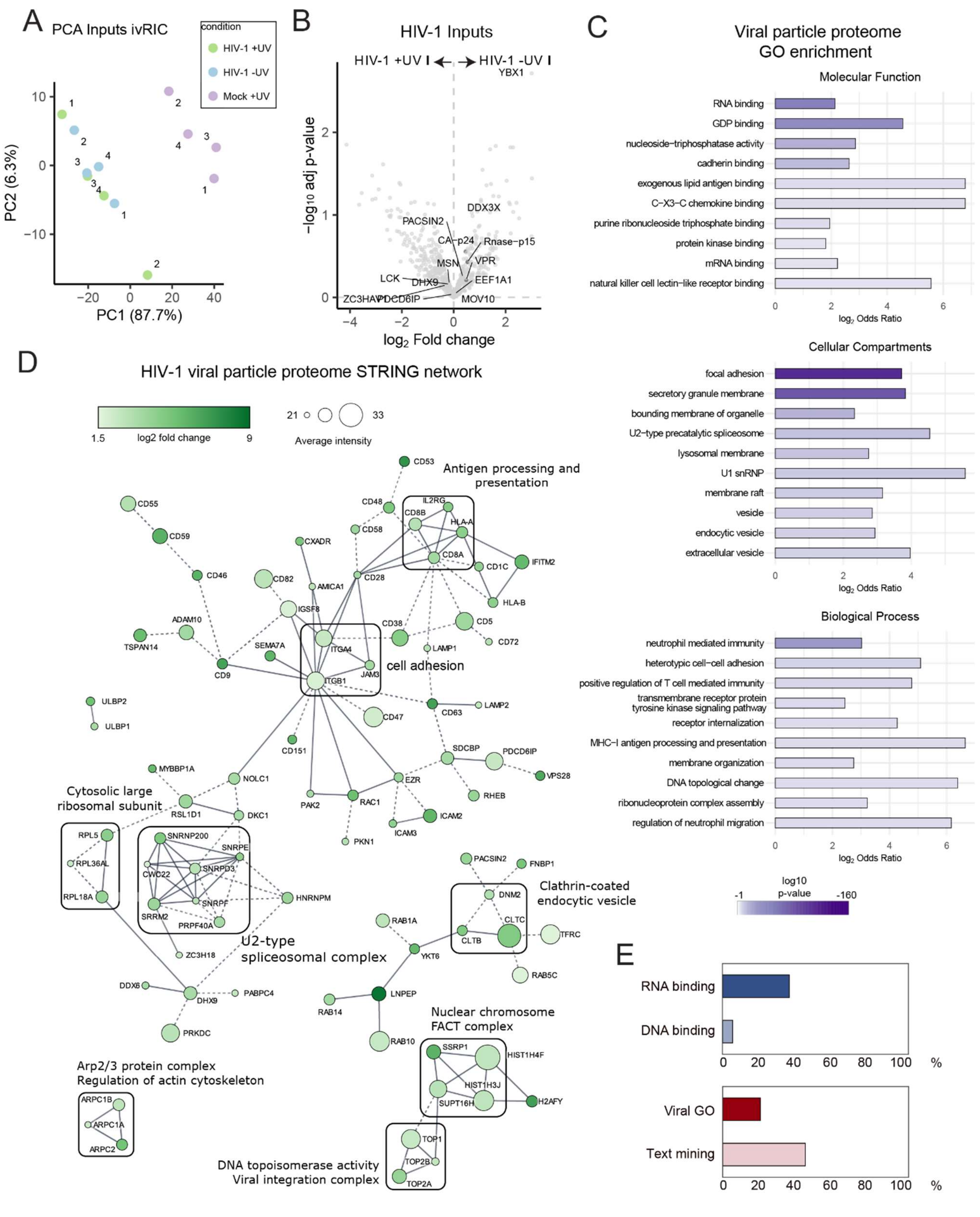


**Figure S3. The proteome of HIV-1 viral particles.** A) Principal component analysis (PCA) of the ivRIC inputs. B) Volcano plot comparing the proteome of viral particles (ivRIC inputs) in UV irradiated and non-irradiated samples. Grey dots are non-significantly enriched proteins. C) GO term enrichment analysis of the proteins enriched in HIV-1 infected (viral particles) over mock-infected inputs. D) STRING network analysis of the proteins enriched in HIV-1 particles using Cytoscape. Physical interactions are represented by solid lines and functional interactions by dashed lines. Top GO enriched terms are shown for each cluster. E) Bar plots showing the proportion of proteins in the viral particles (inputs of ivRIC) annotated with RNA- and DNA-binding (GO terms); virus-related (GO terms) and HIV-1-related (text-mining) functions.
