## Supplementary material for "Incorporation of genome-bound cellular proteins into HIV-1 particles regulates viral infection": Figure S4

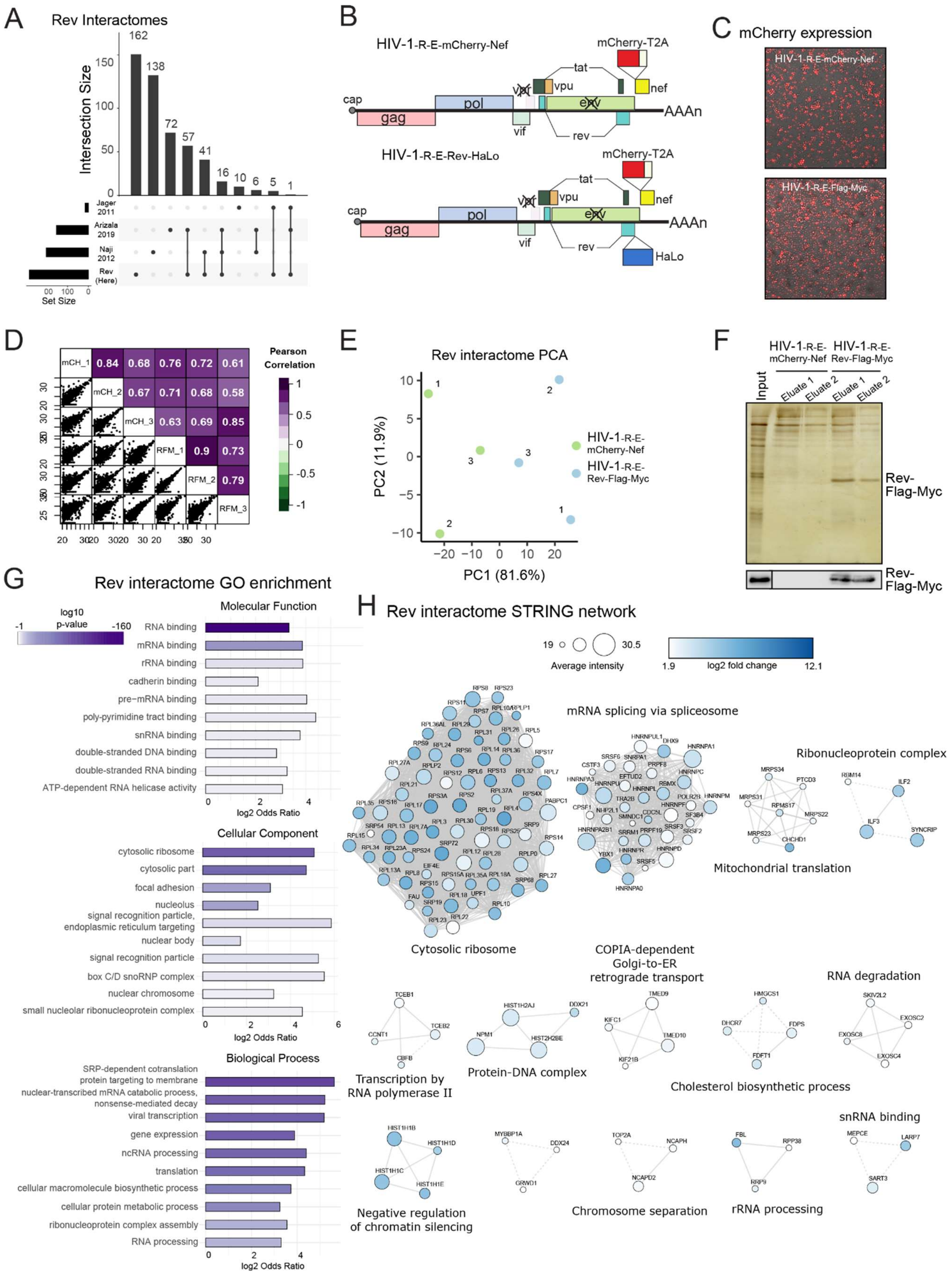


**Figure S4. The proteomic analysis of the Rev interactome.** A) Overlapping of the previously established Rev interactomes with the dataset generated here. B) Schematic representation of HIV-1-R-E-mCherry-Nef and HIV-1-R-E-Rev-HaLo. C) SupT1 cells infected with VSV-G pseudotyped chimeric HIV-1 were observed under a fluorescent microscope and mCherry signal was used as proxy for infection. D) Scatter plots showing the protein intensity and the Pearson correlation between different samples and replicates of the Rev protein-protein interaction experiment. E) PCA of the eluates of the Rev-Flag-Myc IP and the control IPs. F) Silver staining of the Rev-Flag-Myc IP. G) GO enrichment analysis of the Rev interactome.

H) STRING clustered network of the Rev interactome analysed with Cytoscape. Physical interactions are represented by solid lines and functional interactions by dashed lines. Top GO enriched terms are shown for each complex.
