## Supplementary material for "Incorporation of genome-bound cellular proteins into HIV-1 particles regulates viral infection": Figure S5

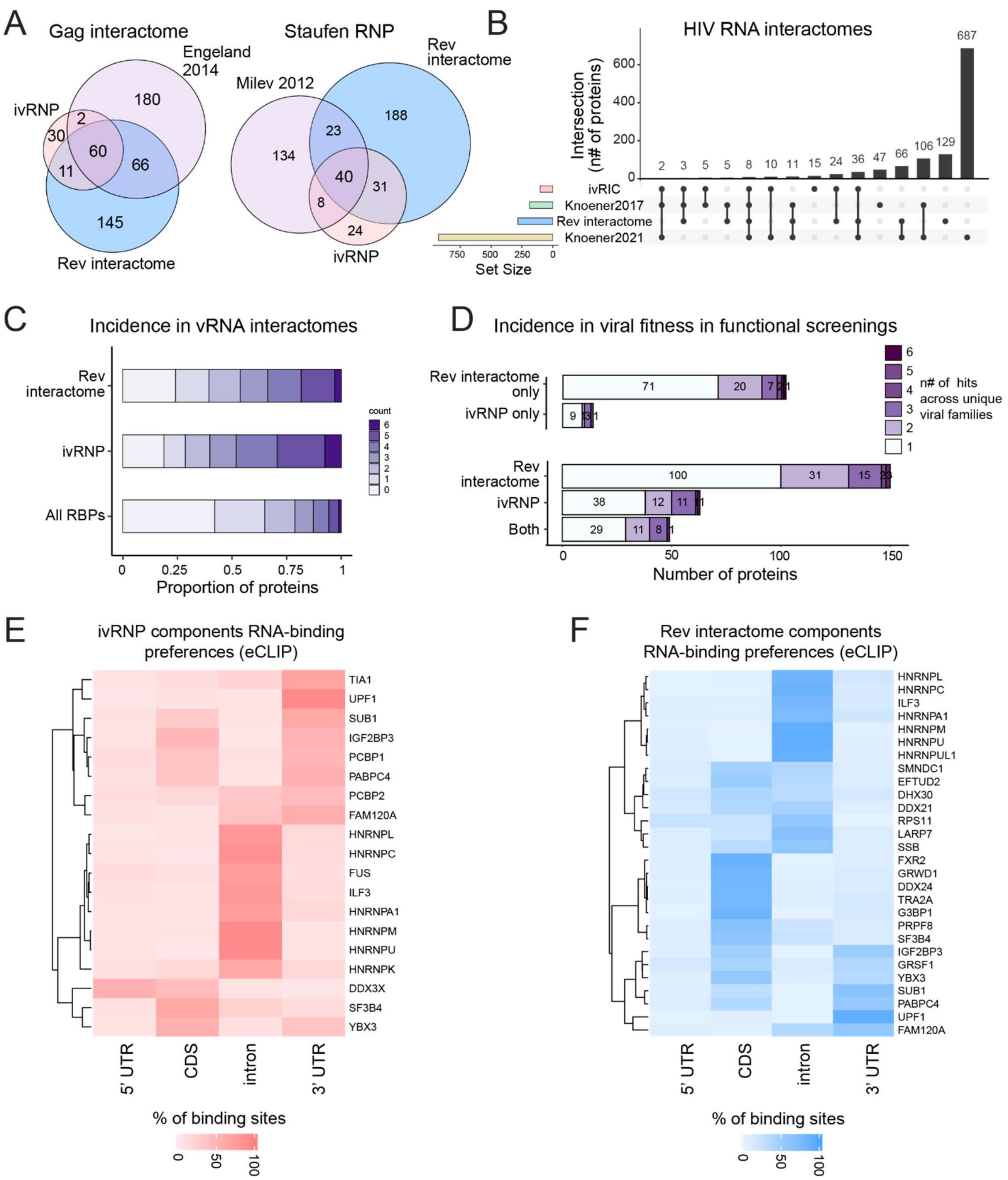


**Figure S5. Characterising the ivRNP and the Rev interactome.** A) Venn diagram showing the overlapping of the ivRNP and the Rev interactome with the previously established Gag[1] and Staufen[2] interactomes. B) Upset plot comparing the ivRNP and Rev interactome to the previously established HIV-1 RNA interactomes[3,4]. C-D) Classification of the ivRBPs and Rev interactors based on their incidence in published viral RNA interactomes (C) or in siRNA and CRISPR/cas9 screenings of virus fitness (D). Colours illustrate the number of different viral families. E-F) Heatmaps showing the incidence of binding sites of ivRBPs (E) and Rev interactors (F) in different regions of the target cellular RNA using the ENCODE eCLIP dataset.
