## Supplementary material for "Incorporation of genome-bound cellular proteins into HIV-1 particles regulates viral infection": Figure S6

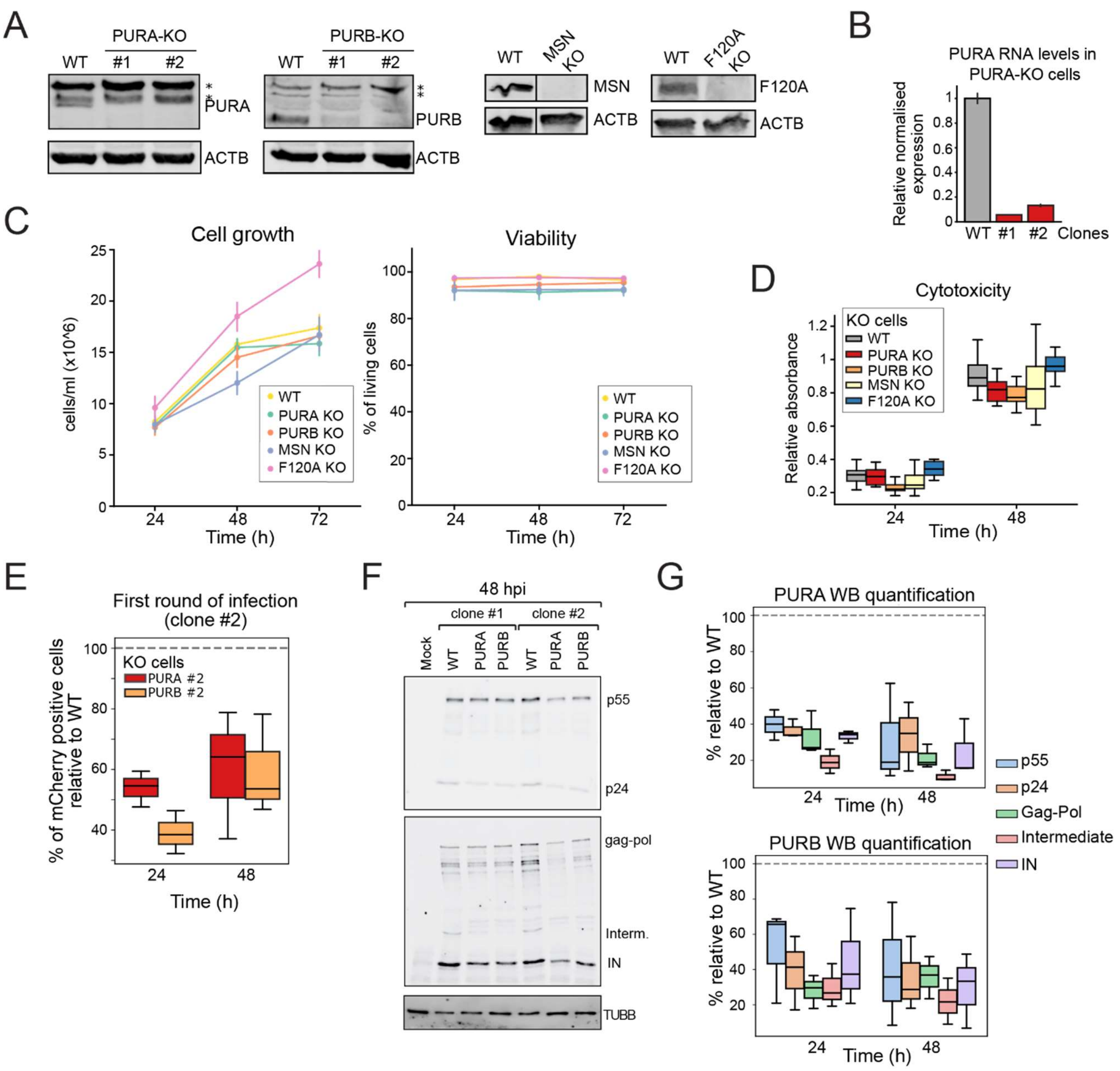


**Figure S6. Establishing KO cell lines for ivRBPs and Rev interactors.** A) Western blot analysis of the different KO cell lines using specific antibodies. B) RT-qPCR analysis of PURA mRNA in PURA KO clones. C) Line plots showing cell proliferation and viability of KO cells. D) Box plot showing cytotoxicity after gene KO. E) Flow cytometry analysis of mCherry positive cells in SupT1 WT, PURA KO and PURB KO clone 2 infected with HIV-1$_{\text{-R-E-mCherry-Nef}}$. F) Western blotting analysis of WT, PURA and PURB KO SupT1 cells infected with HIV-1$_{\text{mCherry-Nef}}$ at 48hpi. G) Box plots showing the Western blot quantification of different HIV-1 proteins from Figures 5C and S6F.
