## Supplementary material for "Incorporation of genome-bound cellular proteins into HIV-1 particles regulates viral infection": Figure S7

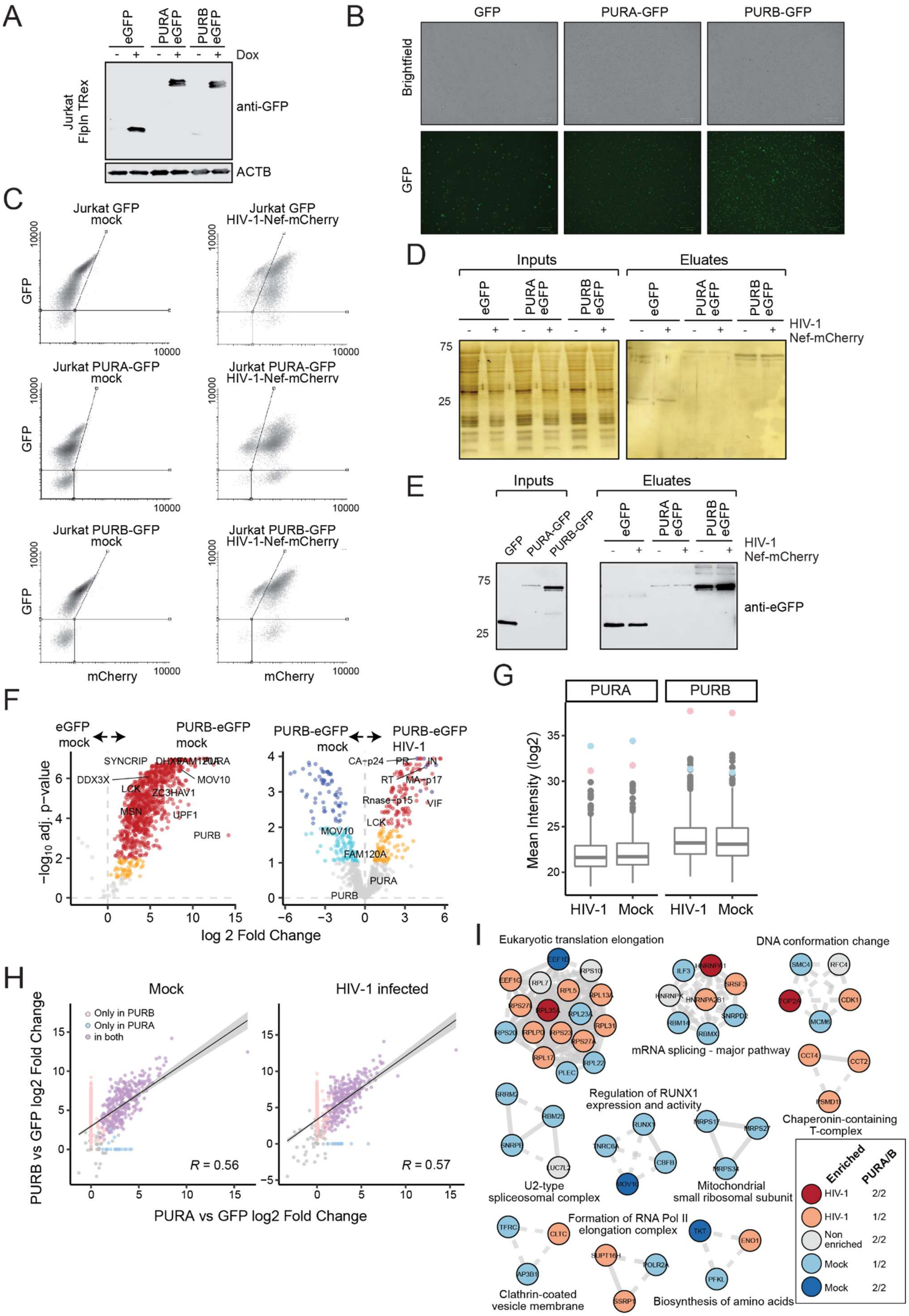


**Figure S7. The protein interactome of PURA and PURB in HIV-1 infected cells.** A) Western blot showing the doxycycline (dox)-inducible expression of PURA-eGFP and PURB-eGFP in Jurkat Flp-In T-Rex. B) Fluorescent protein detection in the different cell lines expressing eGFP. C) Flow cytometry analysis of mock and HIV-1$_{R\text{-}E\text{-}mCherry\text{-}Nef}$ infected cells expressing eGFP-fused proteins at 48 hpi. D-E) Silver staining (D) and Western blot with antibodies against eGFP (E) showing inputs and eluates of the IPs using the cell lysates from panel C and the eGFP nanobody (GFP_Trap). F) Volcano plots showing the enrichment of the PURB-eGFP IP over the eGFP IP (left panel) and PURB-eGFP IP in HIV-1-infected over mock cells (right panel). Red and dark blue dots are proteins enriched with 1% FDR, while orange and cyan dots are proteins enriched with 10% FDR. Grey dots are non-enriched proteins. G) Box plots showing the intensity distribution of proteins in PURA-eGFP and PURB-eGFP IPs. H) Scatter plots comparing the intensity of proteins co-precipitated with PURA-eGFP and PURB-eGFP. I) STRING clustered network of proteins differentially associated to PURA-eGFP or/and PURB-eGFP in mock and HIV-1 infected cells. Top GO terms for each complex are shown. Solid lines represent physical interactions and dashed lines functional interactions.
