## Supplementary material for "Incorporation of genome-bound cellular proteins into HIV-1 particles regulates viral infection": Figure S8

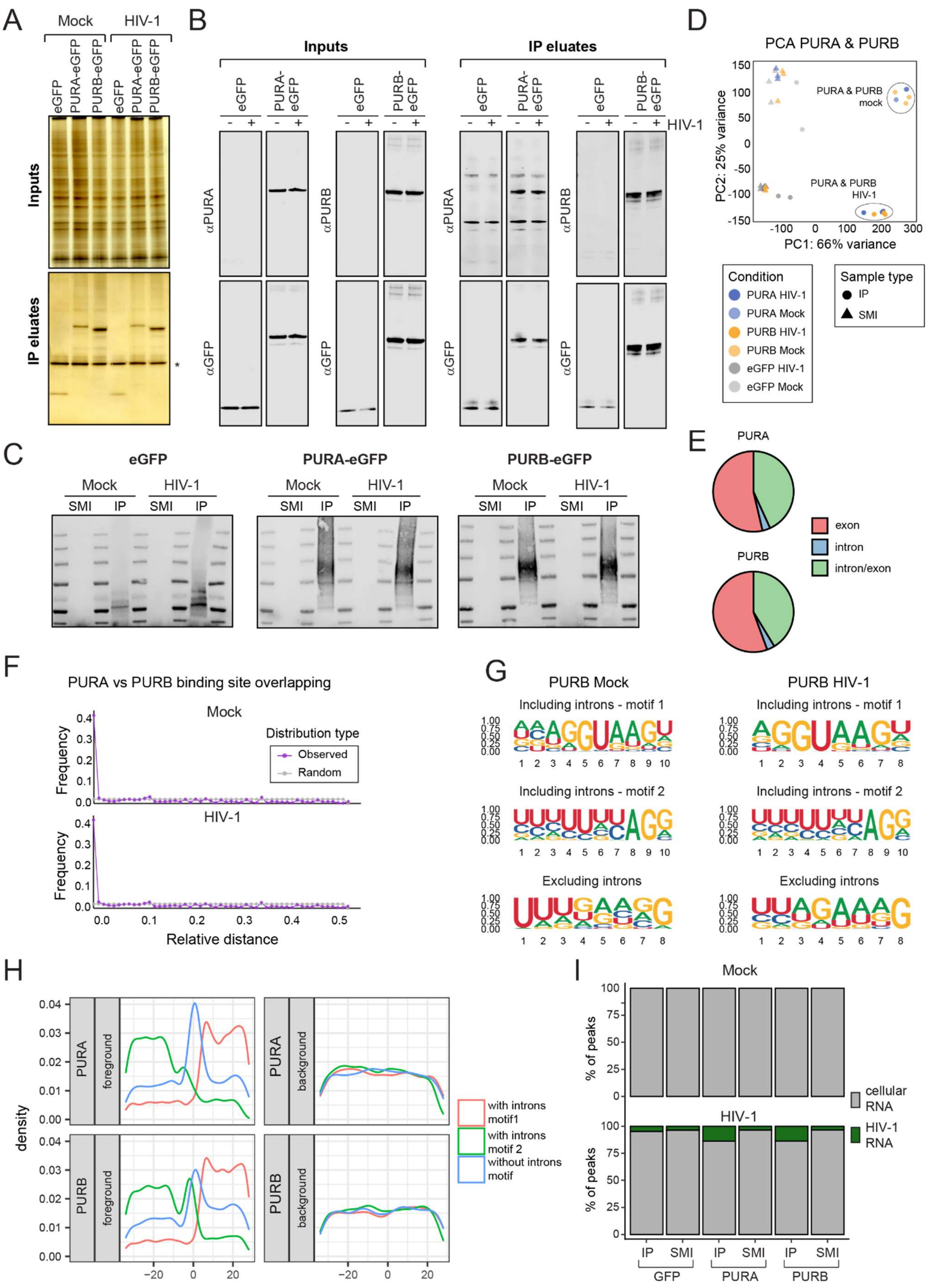

**Figure S8. Analysis of PURA and PURB binding profiles on cellular and viral RNAs.** A-B) Silver staining (A) and Western blot (B) of the inputs and eluates of the IPs for the iCLIP2 experiment (Figure 6) using the cell lysates from Figure S7C and the eGFP_Trap. C) Analysis of the RNA co-purified with the immunoprecipitated protein by ligation of a fluorescent DNA linker at the 3' end and separation by SDS-PAGE. D) PCA of the different iCLIP2 sequencing data. E) Distribution of the binding sites within exons, introns and spanning intron/exon junctions. F) Plot showing the relative distance between PURA-eGFP and PURB-eGFP binding sites. G) Analysis of the sequence motifs recognised by PURB-eGFP using the motif discovery software HOMER and including or excluding introns. H) Density plot showing the distribution of the sequence motifs across the binding site. I) Proportion of iCLIP2 reads mapping to human or HIV-1 genome.
