## Supplementary material for "Incorporation of genome-bound cellular proteins into HIV-1 particles regulates viral infection": Figure S9

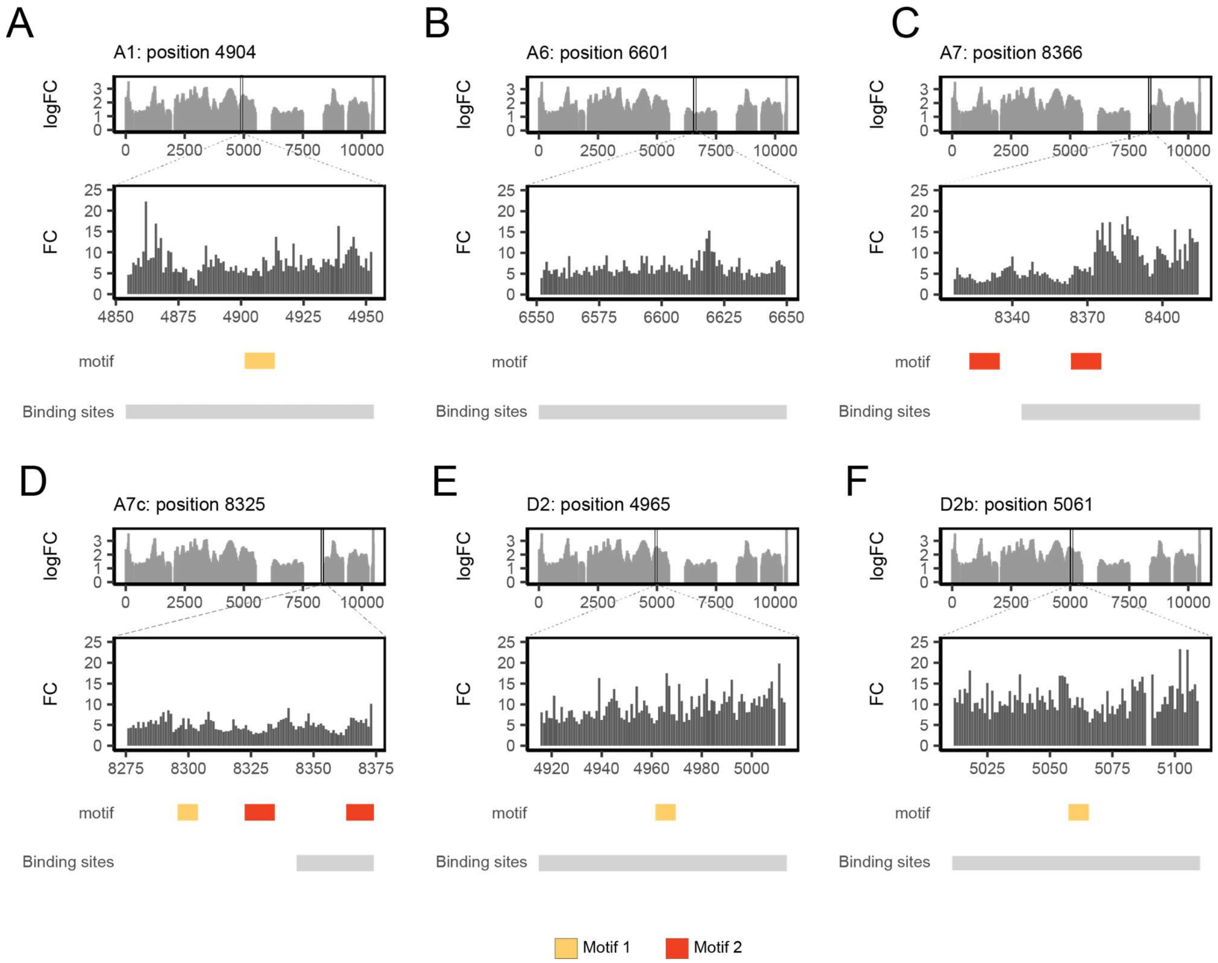


**Figure S9. Analysis of the binding of PURA to the HIV-1 splice sites.** A-F) PURA-eGFP crosslinking intensity around the splice sites on the HIV-1 genome. Only splice sites with significant binding over the SMI control are shown. The position of the two motifs enriched in PURA-eGFP iCLIP2 dataset are indicated (red and yellow). The position of the region with PURA crosslink sites that are significantly enriched over the SMI control (i.e. binding site) are indicated with grey boxes.
